# MIGRATION OF SCHWANN CELL PRECURSORS (SCPS) IS REGULATED BY INTERACTIONS OF N-CADHERIN-MEDIATED ADHESION AND EPHRIN-A2-INDUCED SCP CONTRACTILITY

**DOI:** 10.64898/2026.06.06.730613

**Authors:** Florence K. Roche, Paul C. Letourneau

## Abstract

Schwann cell precursors (SCPs) migrate with peripheral sensory axons from DRGs to target regions. SCP migration involves adhesions to substrata to stabilize advancing cell margins, and contractile forces that pull a SCP forward and break rear adhesions. Ncadherin on axons provides adhesion for SCPs, and ephrin-A2 signaling from axons stimulates SCP contractions to complete SCP translocation. Modulation of these adhesive and contractile forces regulates SCP migratioin during development of peripheral nerves.

## INTRODUCTION

Schwann cells are the supportive glial cells of peripheral nerves (2,11). Every axon, synapse and terminal is enwrapped by Schwann cell processes or a Schwann myelin sheath. Once formed, these Schwann cell wrappings are morphologically stable.

However, when peripheral axons are damaged, Schwann cells re-express active motility and produce growth factors and adhesive molecules. This re-expression of developmental behaviors is critically important in promoting axonal regeneration, nerve repair, and return of function (2,4,23).

During development Schwann cell precursors (SCP) arise from the neural crest and migrate with growing axons into peripheral targets (3, 15, 17–19, 24,39). As SCPs migrate along growing axons, SCPs make frequent contacts with each other and with axons via adhesions mediated by N-cadherin, a member of the cadherin family, which has many important roles in tissue morphogenesis, homeostasis and disease (12,13,35, 38–40). In the absence of N-cadherin function, growth cones *in vitro* do not migrate onto SCPs, SCPs pull away from contacts with growth cones and other SCPs, and SCPs do not migrate in close association with axons (21,22,40). Adhesion molecules NCAM and L1 have also been reported to mediate adhesion between developing Schwann cells and axons (31,34). Migrating SCPs also contact adjacent extracellular matrices (ECM), and express integrins to adhere to laminin, fibronectin and other ECM components (28,30). SCP survival depends on the axon-derived growth factor neuregulin, which keeps migrating SCPs closely associated with growing axons (15). As SCPs migrate, they don’t move ahead of extending axons (3,39). Once all axons reach their targets, SCPs mature to Schwann cells, become less motile, secrete basal lamina components, and begin to enwrap small axons into bundles and ensheath larger axons before myelination (2).

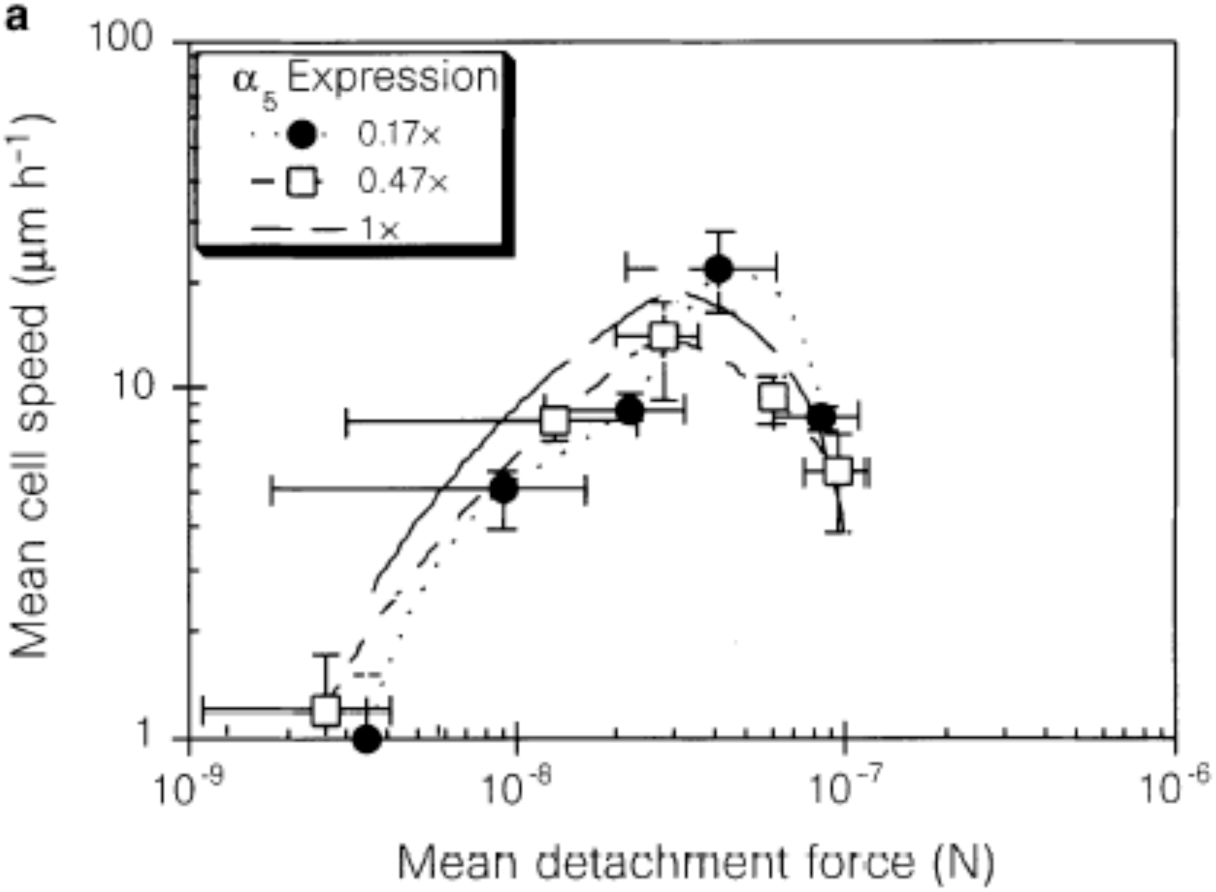

Cell migration is a three-step process that is initiated by cellular extension involving actin filament polymerization at the leading edge. Secondly, the leading margin is anchored by adhesive bonds between surface adhesion molecules and adhesive ligands on the substratum. Thirdly, the motor molecule myosin generates tensions that pull the cell toward its leading attachments and detach the cell posterior from the substratum. Modeling studies show that the speed of cell migration depends in a biphasic manner on cell-substratum attachment strength, with maximum migration occurring at intermediate levels of cell-substratum adhesiveness, which allows formation of new adhesions at the cell front, but also allows rupture of posterior attachments (6,20,26). Graph at left is from Palacek et al. 1997, <u>Nature</u> 383:537-541.

These preliminary studies examine SCP motility and adhesion during a period of active axonal growth and SCP motility.

Ephrins are cell surface-tethered ligands for Eph receptors, which comprise a large family of tyrosine kinase receptors that bind interacting ephrin ligands, and through signaling regulate cell behaviors (5,7,8,9,16,27,29,32,36,43,44). Much ephrin/Eph signaling results in repulsive cell interactions such as in axon guidance, and cell segregation in establishing tissue boundaries and segmentation, and in disease states. Eph A receptors and their Ephrin-A ligands might regulate SCP migration and interactions with axons in two ways. Activation of EphA receptors on SCPs by ephrin-A2 activates the Rho GTPase, which increases myosin contractility in SCPs (32,37).

Secondly, ephrin-A2 activation of EphA increases signaling that can reduce cell adhesion involving both N-cadherin and integrins, weakening SCP-substratum adhesion. Thus, EphA/ephrin-A2 interactions may modulation both the adhesion and migration of SCPs.

We examined the expression of several adhesion molecules, EphA receptors, and ephrin-A ligands on SCPs and axons from E7 and E13 chick DRGs. At E7 in chick embryos large numbers of sensory axons are growing along sensory pathways to peripheral targets (3). At E7 SCPs are dividing and migrating actively in close association with these advancing axons. By E13 most sensory axons have reached target areas. Axonal terminals are branching and forming sensory endings, but new axons are not being added to peripheral nerves. These axons continue to elongate by mechanical "stretched growth" rather than by growth cone migration (33). Until E13 SCPs continue to divide and move to maintain coverage of "stretching" axons. At E13 Schwann cells are maturing, beginning to enwrap axons into bundles, and by E14, myelination is beginning in chick sciatic nerves.

In these preliminary studies we examined the interactions of N-cadherin mediated adhesion and the expression and actions of ephrin-A2 in modulating SCP migration. We found that ephrin-A2 signaling triggers SCP contraction and reduced cell spreading. At E7 ephrin signaling may promote SCP migration, while by E13 ephrin-A2 signaling is very reduced or absent, consistent with limited SCP migration. These preliminary studies were conducted and analyzed in the Letourneau laboratory between mid-2005 and mid-2007. They have not been reported before in an abstract or publication.

## RESULTS

### Culturing DRGs and Schwann cell precursors (SCPs)

We obtained SCPs from two sources in E7 and E13 chick embryos, lumbosacral dorsal root ganglia (DRG) and segments of peripheral nerves distal to lumbar DRGs. When E7 DRG ganglia were explanted onto laminin-coated substrata, SCPs migrated from DRGs in association with hundreds of axons elongating from DRG neurons (Figure 1). These SCPs were identified by their labeling with the HNK antibody (Fig. 2A), a surface carbohydrate epitope on neural crest derivatives, by the 1E8 antibody (D), which labels the SC lineage, and by expression of the p75 neurotrophin receptor. SCPs from DRGs and peripheral nerves also expressed cell adhesion molecules NCAM (B), NrCAM, and N-cadherin (C), but expression of L1 (E) is weak on E7 SCPs. E7 SCPs were also labeled by antibodies against ephrin A receptors, EphA3 and EphA4 (Fig. 2F).

**Figure 1.**
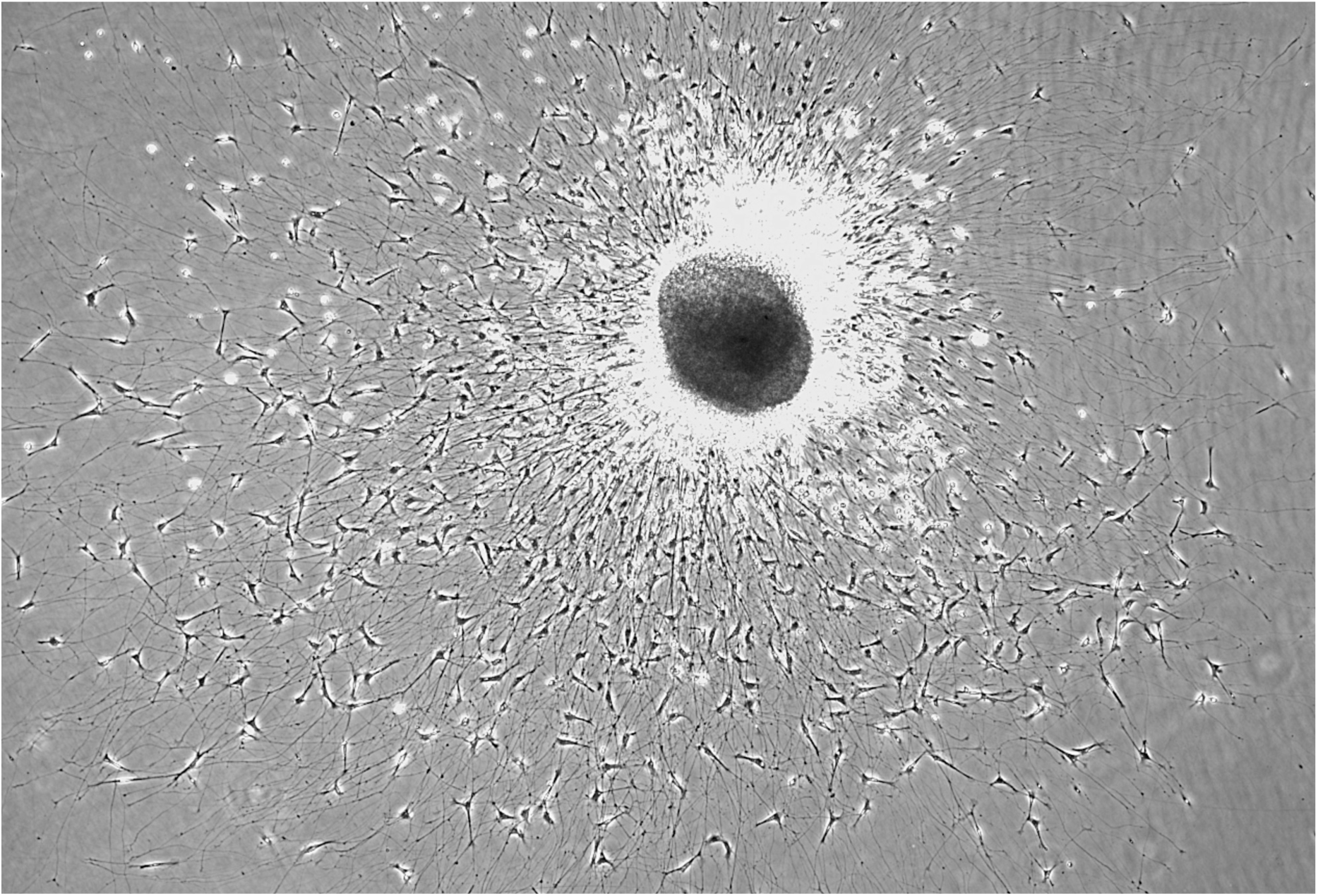
An E7 DRG explant cultured on laminin in high [Ca++] medium. Hundreds of SCPs migrate from the explant along axons and the laminin substratum. 10X.

**Figure 2.**
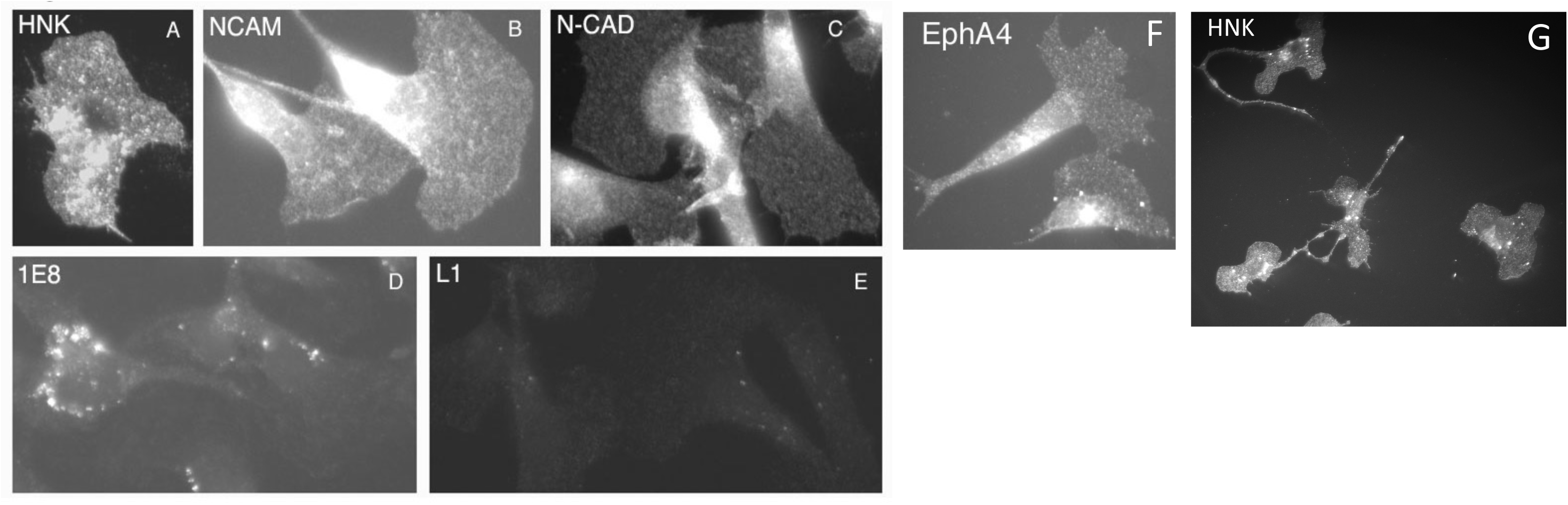
A-F E7 SCPs labeled with antibodies against HNK (A), NCAM (B), N-cadherin (C), 1E8 avian Schwann cell marker (D), L1 (E), EphA4 receptor (F). 2G an E13 SCP labeled for HNK antigen. 63X.

We also prepared cultures of dissociated E7 and E13 chick peripheral nerve explants. E13 explants contain a mix of SCPs, immature Schwann cells, and perineural cells. SCPs were recognized by double labeling with the HNK antibody and antibodies against Schwann cell markers (Fig. 2G). SCPs were labeled with antibodies against adhesion molecules, L1, NCAM, and N-cadherin, the p75 neurotrophin receptor and with the 1E8 antibody. E13 SCPs were weakly labeled with antibodies against ephrin-A receptors, EphA3 or EphA4.

### Expression of cell adhesion molecules, EphA4 and ephrin-A2 on axons and SCPs

We labeled axons extended from explanted E7 and E13 DRGs with antibodies against cell adhesion molecules, L1, NCAM, and N-cadherin. These axons were also labeled by antibodies against ephrin-A2 and –A5, although labeling was very weak for ephrin-A5.

In order to compare fluorescence staining for these surface components on E7 and E13 DRG axons and on E7 and E13 SCPs, cultures from E7 and E13 embryos were prepared, fixed, stained, and photographed together identically. The mean intensity of immunofluorescence was determined in equivalent regions of a random sample of E7 and E13 axons and SCPs.

N-cadherin, NCAM, and L1 were present on E7 and E13 axons (Fig. 3). L1 staining appeared similar on E7 and E13 axons, NCAM expression appeared stronger on E7 axons, and N-cadherin expression was stronger on E13 axons. Ephrin-A2 was expressed on E7 DRG axons, and at much lower levels on E13 axons (Fig. 4). This decreased expression of ephrin-A2 on axons of older embryos is like *in vivo* finding of Iwamasa et al (16).

**Figure 3.**
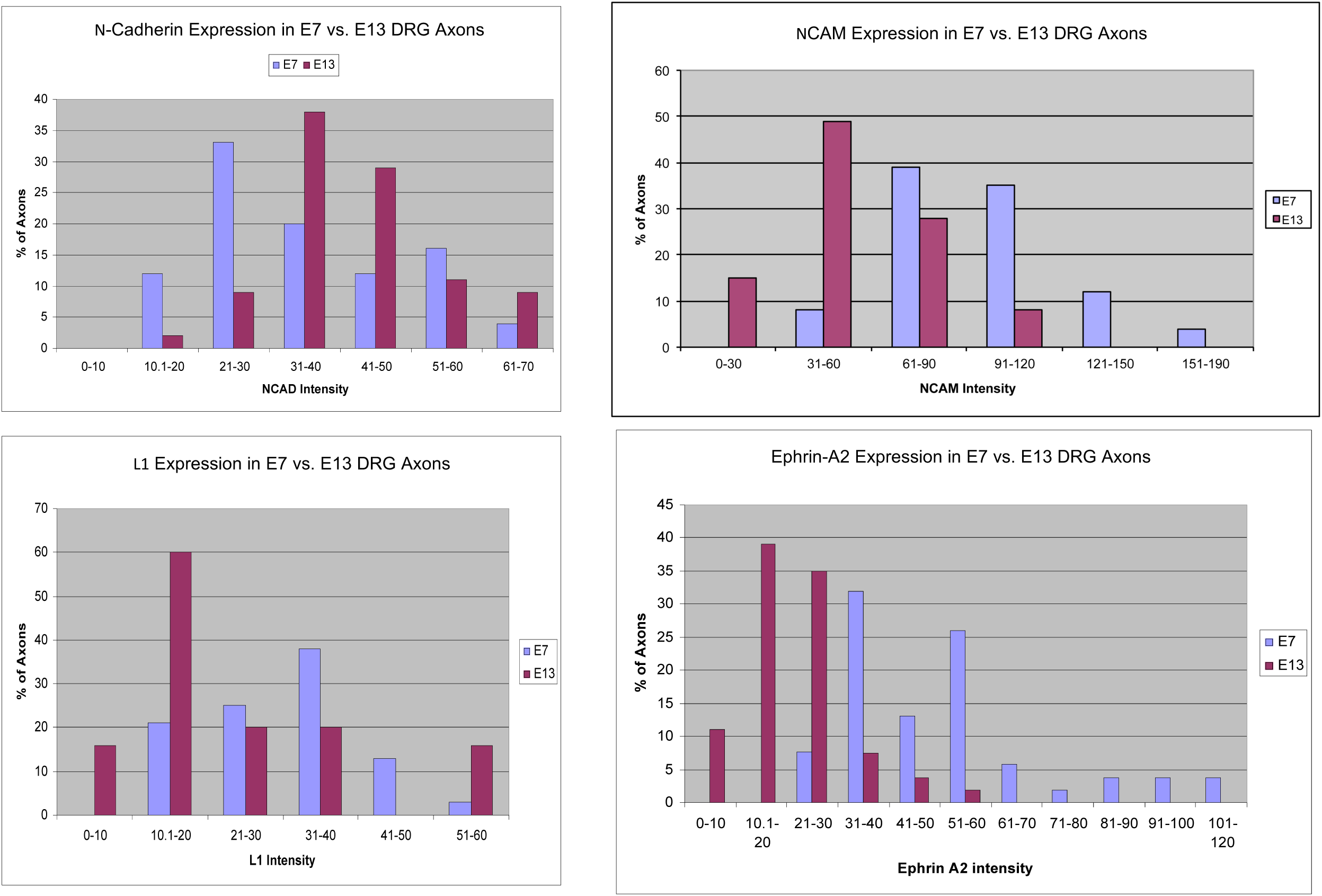
Quantification of immunocytochemical labeling of E7 and E13 DRG axons for expression of N-cadherin, NCAM, L1 and ephrin-A2. Each graph represents measurements of at least 50 axons from E7 and E13 DRG explants.

**Figure 4.**
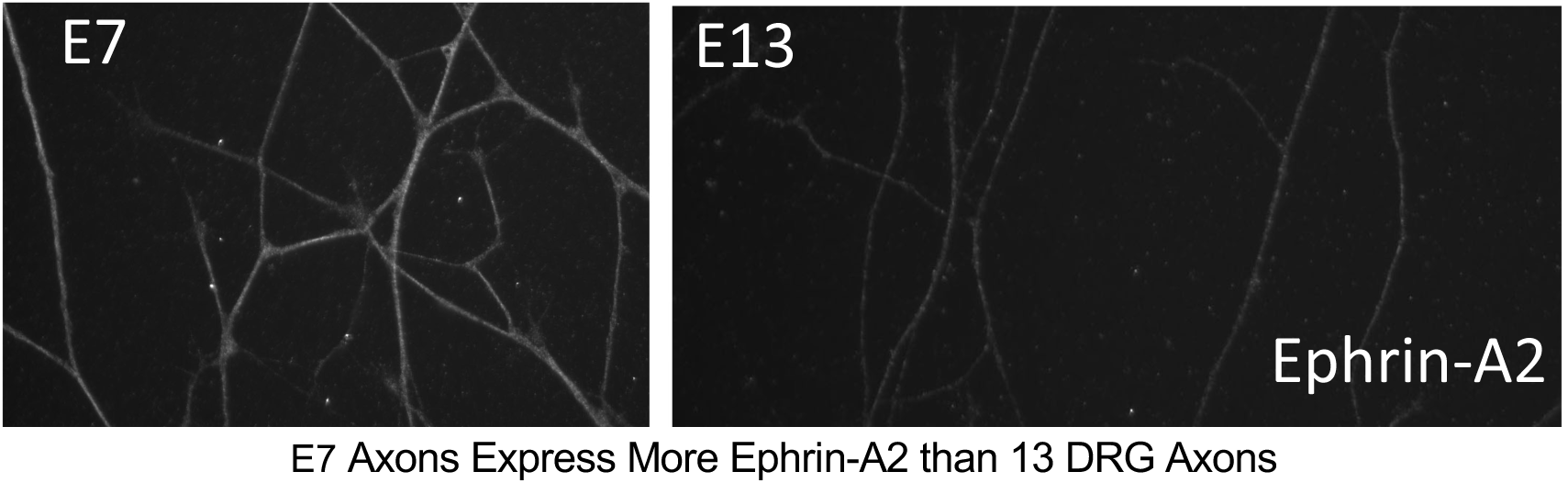
E7 DRG axons label much stronger for ephrinA2 ligand than E13 axons do.

E7 and E13 SCPs from peripheral nerve explants were stained for these same adhesion molecules (Figure 5). In addition, we measured immunofluorescence of ephrin-A2 and EphA4, because labeling for these two proteins on axons and SCP was much stronger than for ephrin-A5, EphA3 or EphA5.

**Figure 5.**
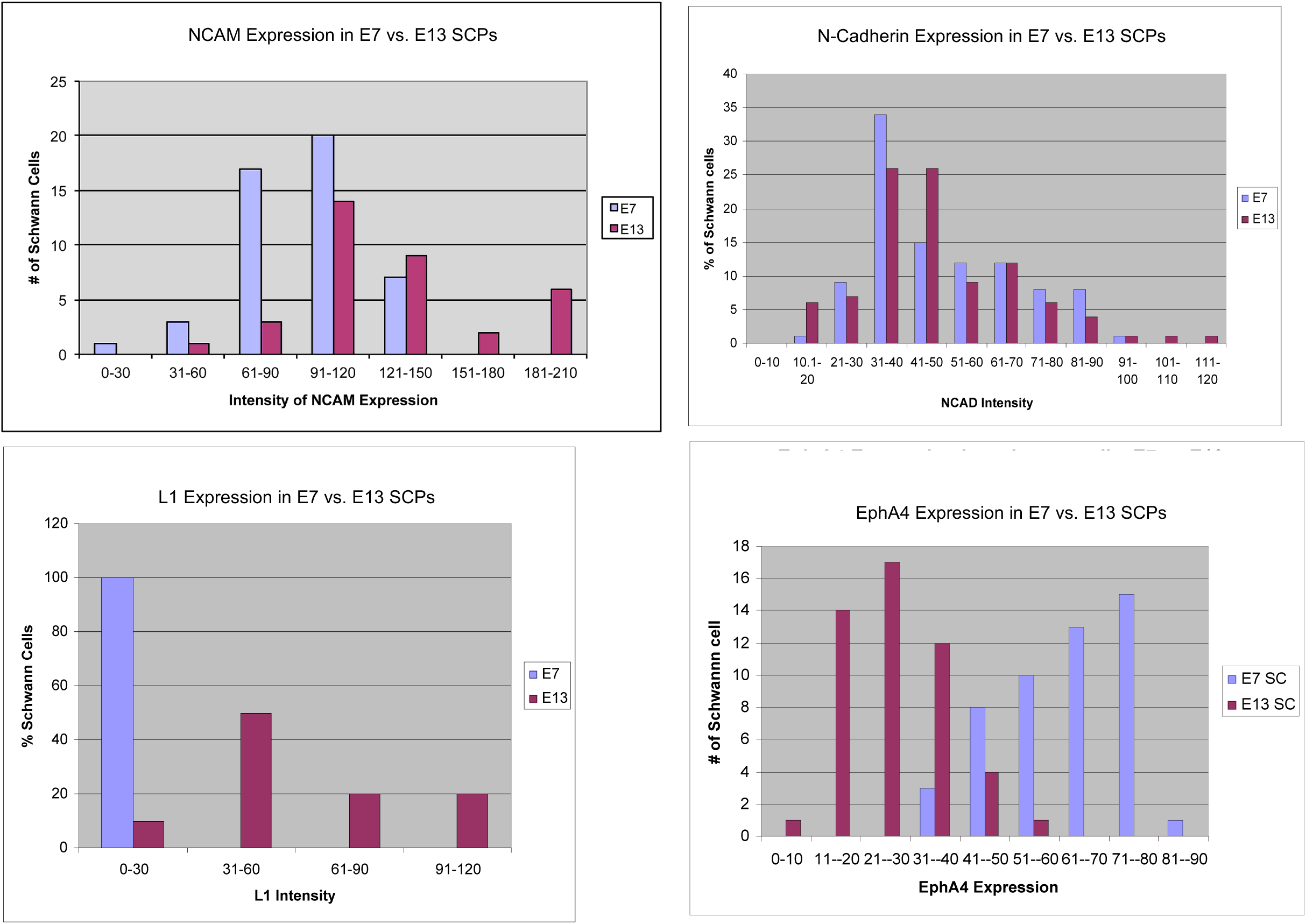
Quantification of immunocytochemical labeling of E7 and E13 DRG SCPs for expression of N-cadherin, NCAM, L1 and EphA4. Each graph represents measurements of at least 50 SPCS from E7 and E13 DRG explants.

The neural adhesion molecules NCAM and N-cadherin were expressed on both E7 SCP and E13 SCPs. NCAM staining appeared brighter on E13 than E7 SCPs. E 13 SCPs expressed L1, but L1 expression was absent or very low on E7 SCPs. E7 SCPs expressed EphA4, while E13 SCPs showed much lower expression of EphA4. Both E7 and E13 SCPs expressed low levels of ephrin-A2 with perhaps higher levels on E7 SCPs.

In summary, the adhesion molecules, L1, N-cadherin, and NCAM were expressed on E7 and E13 axons, while the repulsive ligand ephrin-A2 was present on E7 DRG axons, but not on E13 axons. For SCPs, both E7 and E13 SCPs expressed N-cadherin and NCAM, but E7 SCPs lacked L1 expression. E7 SCPs expressed the ephrin-A2 receptor EphA4 and ephrin-A2 at higher levels than expressed on E13 SCPs. Thus, at E7 expression of adhesion molecules, ephrin-A2 and EphA4, on axons and SCPs would be consistent with both adhesive and perhaps repulsive interactions between axons and SCPs at E7, while at E13 adhesive interactions are favored.

### Motility of SCPs from nerve explants on laminin

On laminin-coated substrata SCPs migrated away from E7 sciatic nerve segments. The SCPs were well spread and migrated as rapidly at >100 µm/hour (Video 1). Protrusion and adhesion of the leading cell margin advanced the cell front, while contractility at the cell rear retracted the trailing cell processes (Fig. 6a; Video 2).

**Figure 6a.**
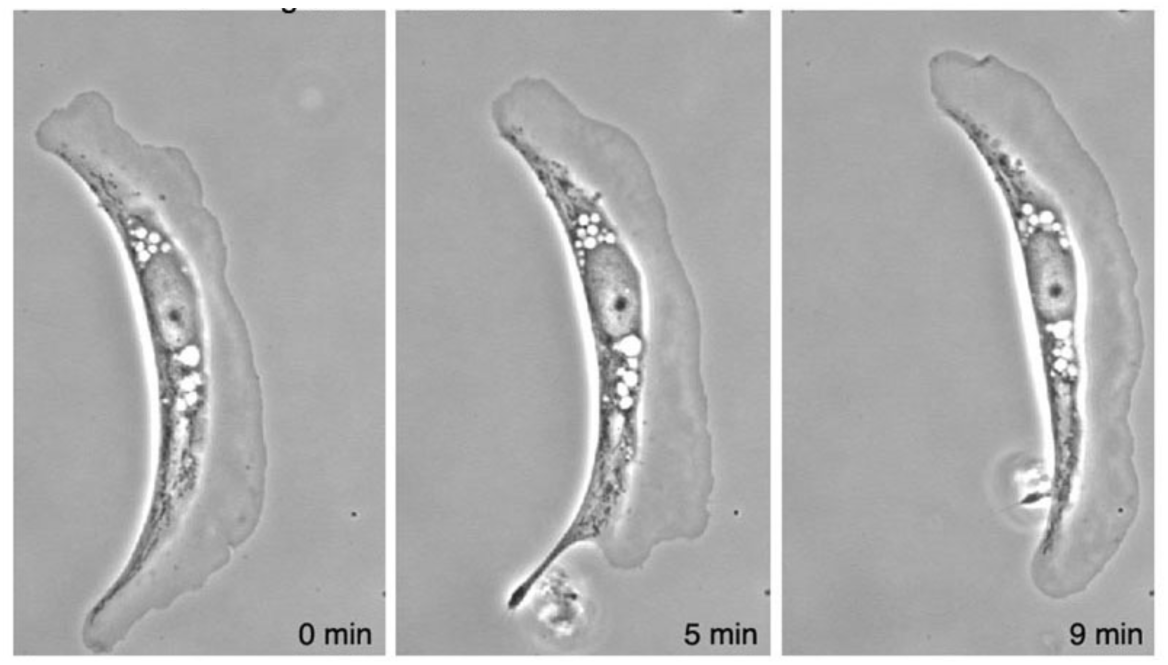
Frames of a migrating E7 SCP. The cell moves with a broad motile leading lamella and the tail is retracted within 4 minutes. 63X.

**Figure 6b.**
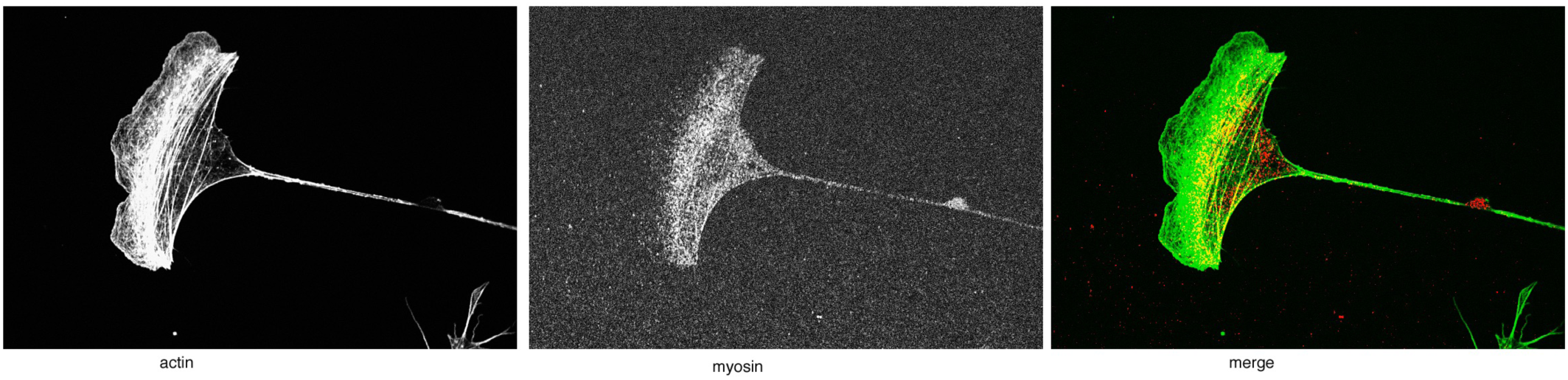
A migrating E7 SCP labeled with fluorescent phalloidin for actin (green) and anti-Myosin II (red). Myosin II-mediated contractility organizes actin filaments into bundles (green). 63X.

**Figure 6c.**
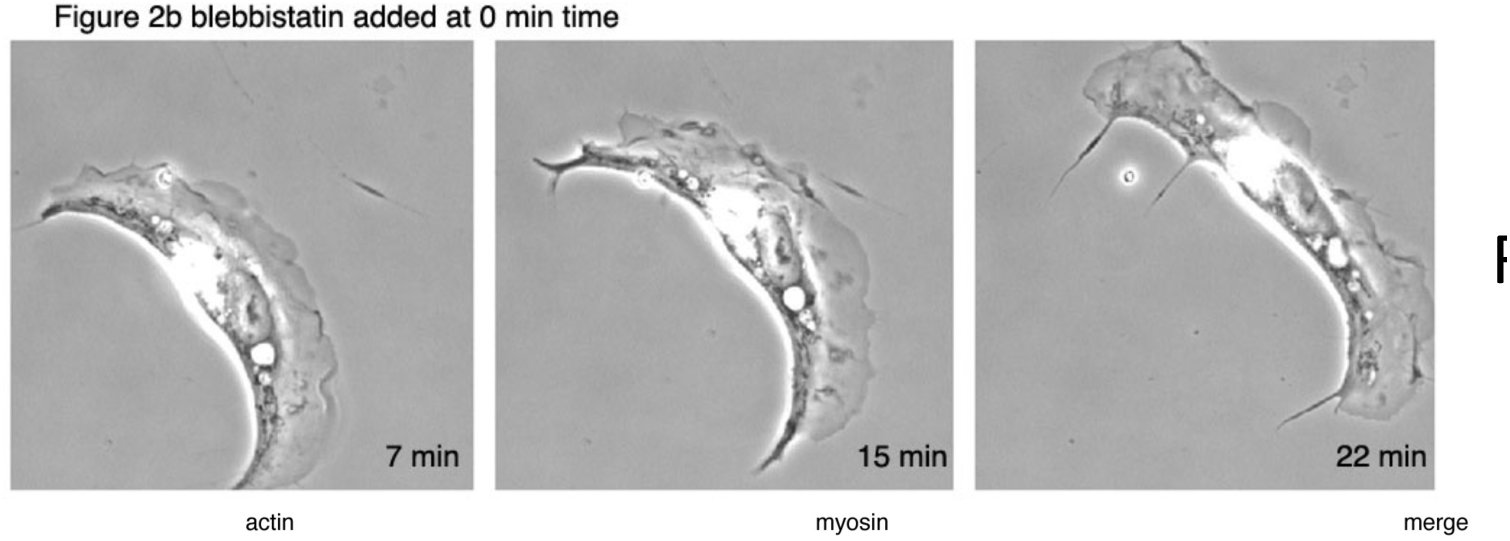
The same SCP as shown in 6a after addition of the myosin II inhibitor blebbistatin. Notice how the cell leading lamella has become broader and the cell tail is not retracted. 63X.

**Figure 6d.**
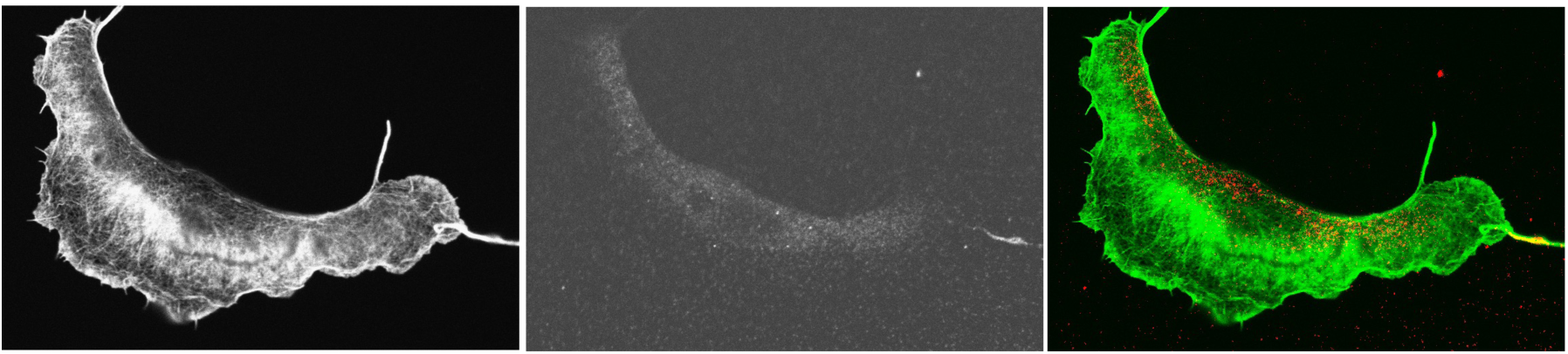
A migrating E7 SCP treated with 20 min with blebbistatin. Actin is labeled with fluorescent phalloidin (green) and myosin II is labeled with anti-myosin II (red). Notice that the inactive myosin (red) is not localized with the actin filaments and actin filament bundles are not present. 63X.

Dense networks of short dynamic actin filaments filled the leading lamellae, and in the cell posterior actin filaments were organized into bundles by myosin II that generated contractile forces to retract the cell tail (Fig. 6b). When migrating SCPs were treated with the inhibitor of non-muscle myosin II, blebbistatin, the migration rate decreased by 50%. After adding blebbistatin, the leading lamella often broadened and the cells lost the polarity of a broad front and a narrower extended rear (Fig 6c). In the blebbistatin-treated SCPs posterior actin filament bundles disappeared (Fig. 6d), and cell tails were not retracted (Fig. 6c; Video 3). Compare images of the rapid tail retraction in Fig. 6a to the absence of retraction in Fig. 6c, seen in videos (Videos 2,3) of the same cell before and after the addition of blebbistatin.

### Migration of SCPs from nerve explants on n-cadherin and on L1

When E7 sciatic nerve explants were cultured on substrata coated with N-cadherin, SCPs also migrated away from the explants, although not as rapidly as on laminin. The adhesive function of N-cadherin is regulated by external [Ca++] (1), such that N-cadherin function ceases below 0.3 mM [Ca++], while maximum adhesive function occurs at 0.8 mM extracellular [Ca++]. Thus, we examined the role of N-cadherin mediated adhesion in SCP migration from peripheral nerve explants. When sciatic nerve segments were explanted onto a laminin or an N-cadherin substratum in medium with 0.3 mM [Ca++] SCPs moved away from the explant as individuals, while in media with 1 mM [Ca++], SCPs migrated away from explants while maintaining contact with neighboring cells, almost as a sheet of cells (Figure 7). E7 Sciatic nerve segments do not attach to an L1-coated substratum, reflecting the lack of L1 expression by E7 SCPs.

**Figure 7.**
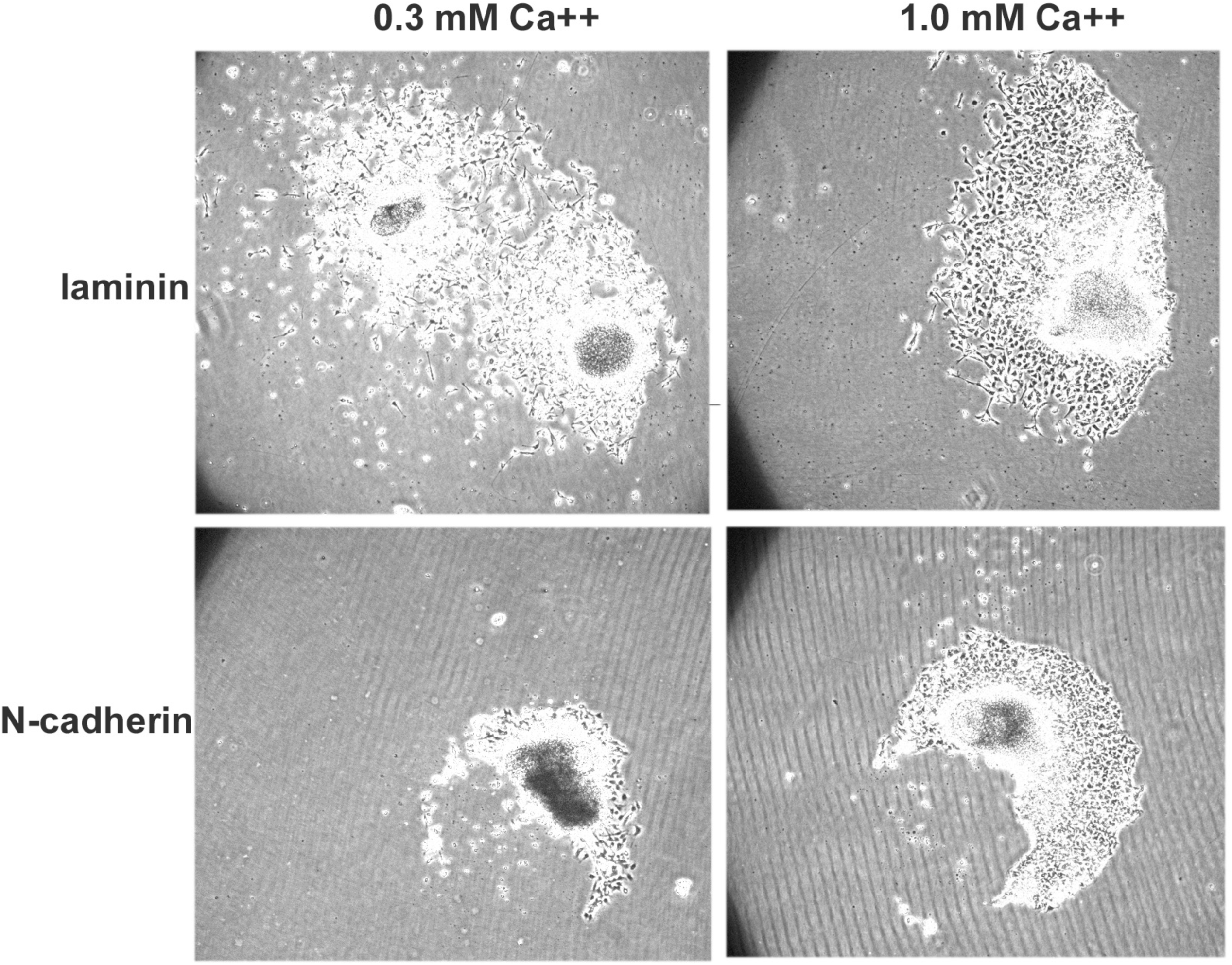
Segments of E7 peripheral nerves cultured overnight on a laminin or N-cadherin substratum. In the presence of 0.3 mM Ca++, N-cadherin mediated adhesion is low, but in the presence of 1.0 mM Ca++ N-cadherin is fully functional and the SCPs maintain cell-adhesions as they migrate away from the explant as a cell sheet. 10X.

### Ephrin-A2 regulates SCP motility and adhesion

The expression of EphA4 on E7 SCPs suggests that SCP may respond to ephrin-As. We directly demonstrated this by visualizing responses of SCPs to contact with beads coated with ephrin-A2-Fc or N-cadherin-Fc. When the leading lamella of a SCP touched an N-cadherin bead, the bead adhered and was transported back on the leading lamella, as the lamella and the cell continued to advance (Video 4). This retrograde transport of ligand-coated beads occurred when cell surface receptors bind to the bead and link to the actin cytoskeleton. SCP responses to ephrin-A2 beads were different.

When Ephrin-A2 beads contacted the edge of a lamellipodium, they adhered, and the lamellar region where the ephrin-A2 bead was bound contracted (Fig 8; Video 5). In addition, sometimes, when an ephrin-A2 bead was picked up by the leading lamella, that region stopped protrusion, while the remainder of the lamella advanced. When ephrin-A2 beads landed on SCP cell tails, the tails rapidly retracted (Fig. 9; Video 6). These contractile events were inhibited by the myosin II inhibitor blebbistatin or the Rho kinase inhibitor, H1152. These results showed that SCP are sensitive to ephrin-A2, and SCP responses to ephrin-A2 beads indicated that signaling from ephrin-A2 may promote local contractility and locally inhibit actin polymerization.

**Figure 8.**
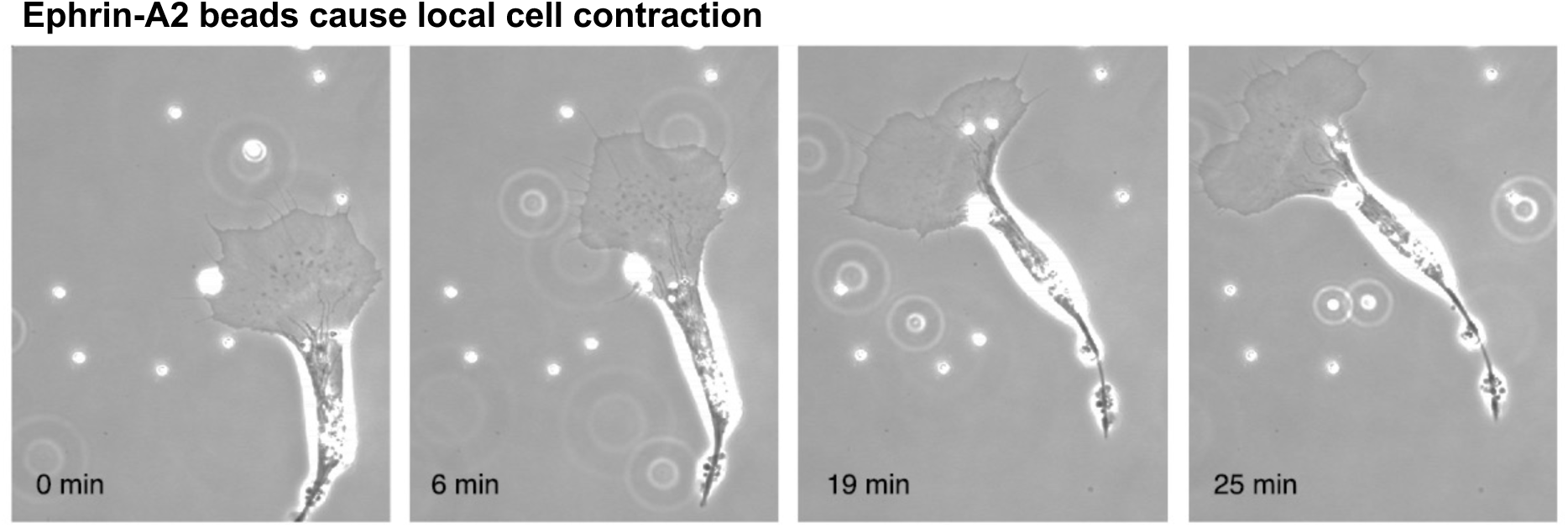
An E7 SCP encounters ephrin-A2 coated beads. The beads adhere to leading lamella and stimulate local contraction of the leading lamella. 63X.

**Figure 9.**
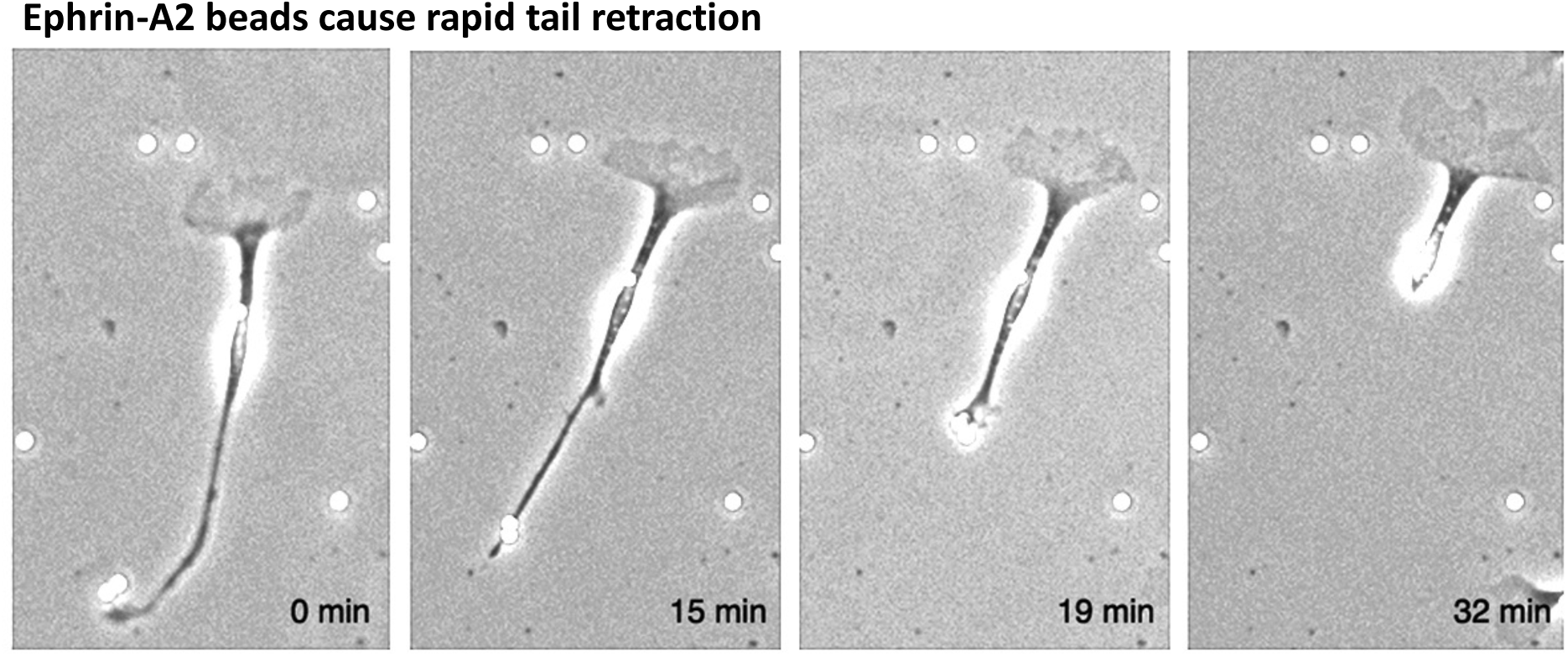
The tail of an E7 SCP touches an ephrin-A2 bead, and the tail is retracted into the cell body. 63X.

We analyzed time lapse videos starting when an SCP leading lamella picked up a bead and then every 30 sec measuring the distance from the cell nucleus to the front edge of the leading lamella along a line passing through the bead (on-axis) or along an adjacent line from the nucleus to the leading lamella, but away from the site of bead contact (off-axis). For cells that contacted N-cadherin beads, the lengths of the on-axis and off-axis lines remained equal (Fig. 10), indicating that the N-cadherin beads did not affect the advance of the leading lamella. However, for cells that contacted ephrin-A2 beads, the on-axis line shortened over time as the off-axis line retained its initial length as the cell advanced (Fig. 11). This response suggested that contact with ephrin-A2 beads caused local contraction and/or cessation of lamellar protrusion.

**Figure 10.**
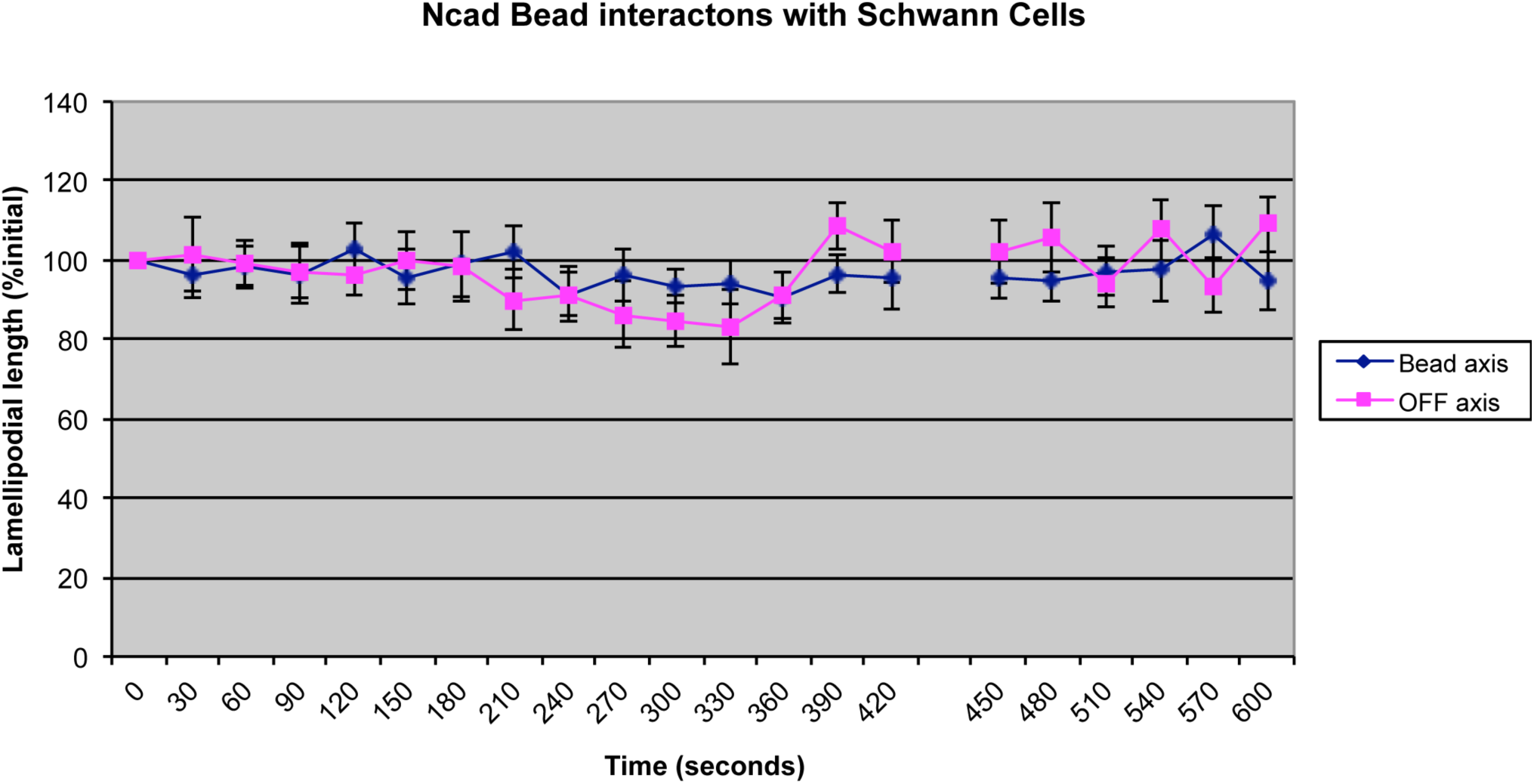
Graph shows that the adhesion of an N-cadherin-coated bead to the front of an E7 SCP does not cause local contraction or cessation of the leading lamella. Each point represents measurements taken from at least 20 video records of SCPs.

**Figure 11.**
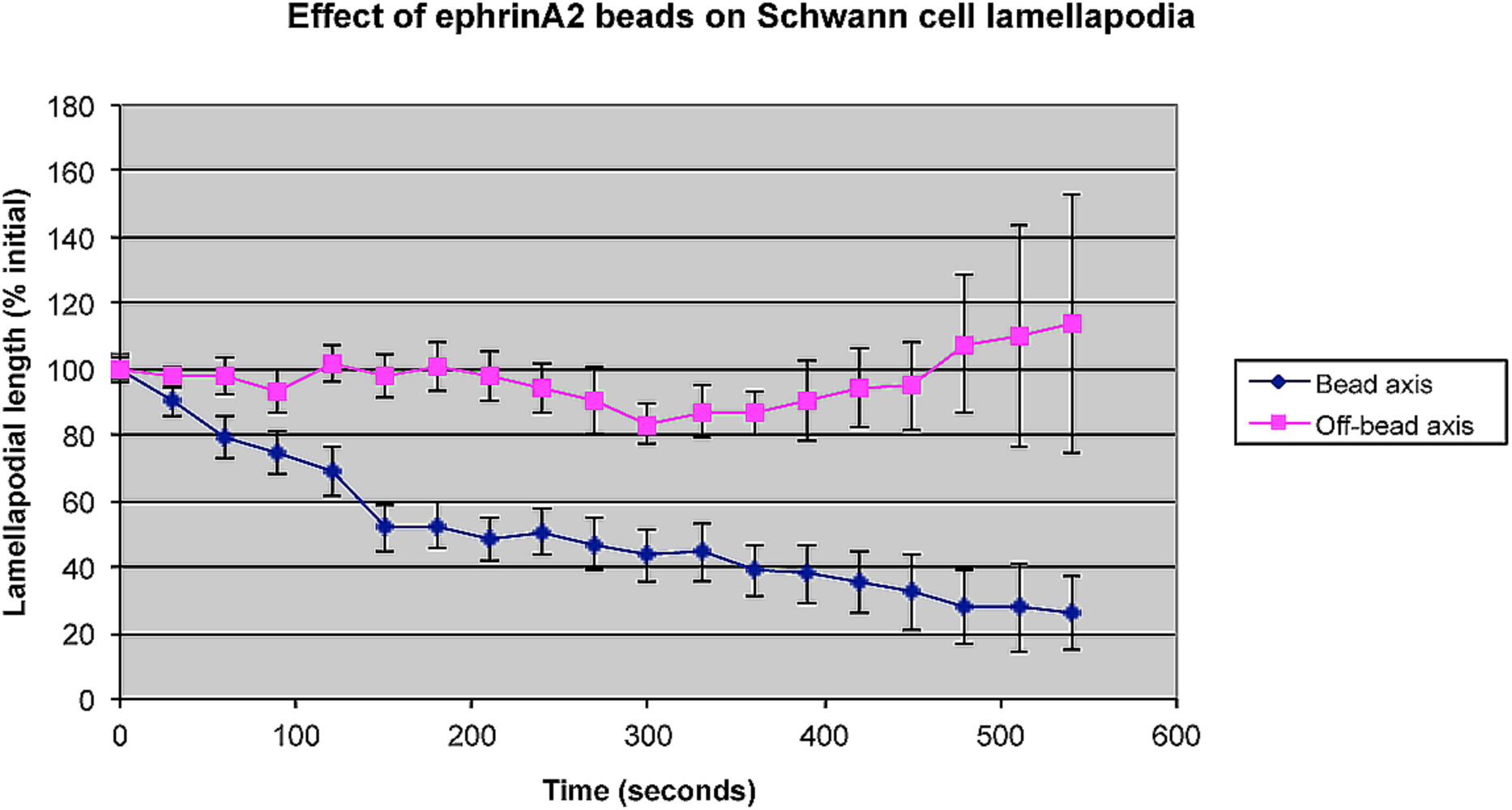
Graph shows that the adhesion of an ephrin-A2 -coated bead to the front of an E7 SCP causes local contraction or cessation of the leading lamella. Each point represents measurements taken from at least 20 video records of SCPs.

We also conducted experiments in which we increased SCP interactions with ephrin-A2 by coating substrata with either 2 ug/ml laminin or 2 ug/ml N-cadherin mixed with 2 ug/ml of an ephrinA2-Fc construct or a control Fc. Figure 12 shows explants of E7 peripheral nerve segments plated for 7 hr. on substrata of laminin-Fc or laminin-ephrin-A2-Fc. By 7 hr. many SCPs had migrated from the explants of laminin-Fc but very few had migrated onto the laminin-ephrin-A2-Fc substratum. We prepared suspensions of E7 SCPs, plated them onto these substrata for 60 minutes, and then photographed sample microscope fields to determine the percent of SCPs that had lamellae and had flattened onto the substratum vs. rounded SCPs without lamellae. The percent of SCPs that had flattened and formed lamellae on laminin or N-cadherin was substantially reduced on substrata which also included ephrinA2-Fc (Fig. 13,14). The src family kinase, Abelson kinase, mediates the repulsive effects of ephrins on nerve growth cone collapse and axon retraction (14). When we included the Abelson kinase inhibitor STI571 in the culture medium of dissociated SCPs, this blocked the inhibition of SCPs flattening and lamellipodial formation on substrata that contained the ephrinA2-Fc construct (Figs. 13, 14, 16). This result suggested that the effects of ephrin-A2 on SCP adhesion and flattening was mediated by Abelson kinase. Murai et al. (25) made a peptide (KYL peptide) that binds tightly and selectivity to the ephrin-A binding site of EphA4 to competitively inhibit ephrin-A2 activation of EphA4. Dr. Elena Pasquale sent us this peptide and a control peptide. When we added this blocking peptide to media with SCPs plated on a laminin-ephrin-A2-Fc substratum, SCP spreading was not inhibited by the presence of ephrin-A2 (Fig. 15).

**Figure 12.**
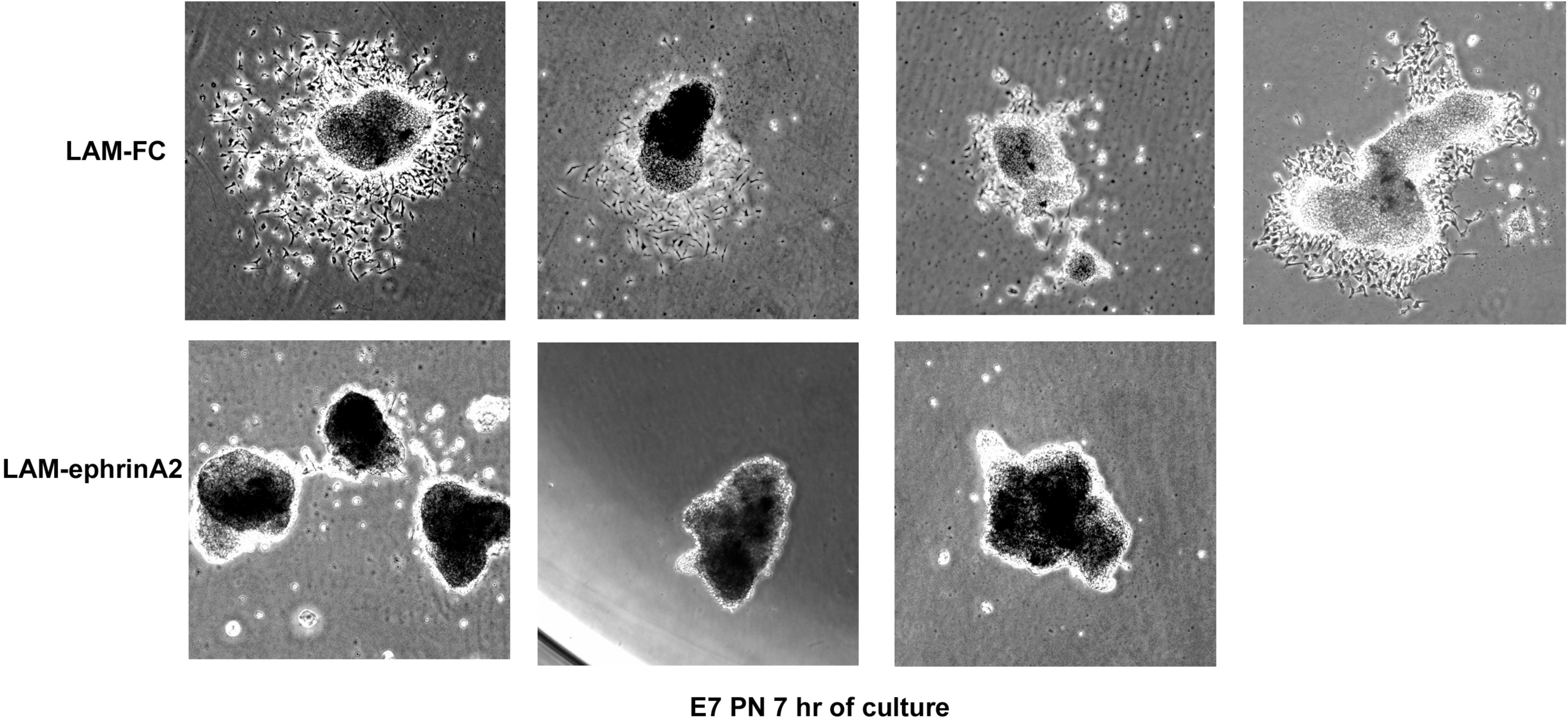
Segments of E7 SCPs were explanted onto substrata coated with laminin-Fc or a laminin-ephrin-A2-Fc mixture. After 7 h *in vitro* explants on laminin-Fc adhere and SCPs can be seen migrating from the explant. However, on laminin-ephrin-A2-Fc explants are poorly attached and few SCPs have migrated from the explants.

**Figure 13.**
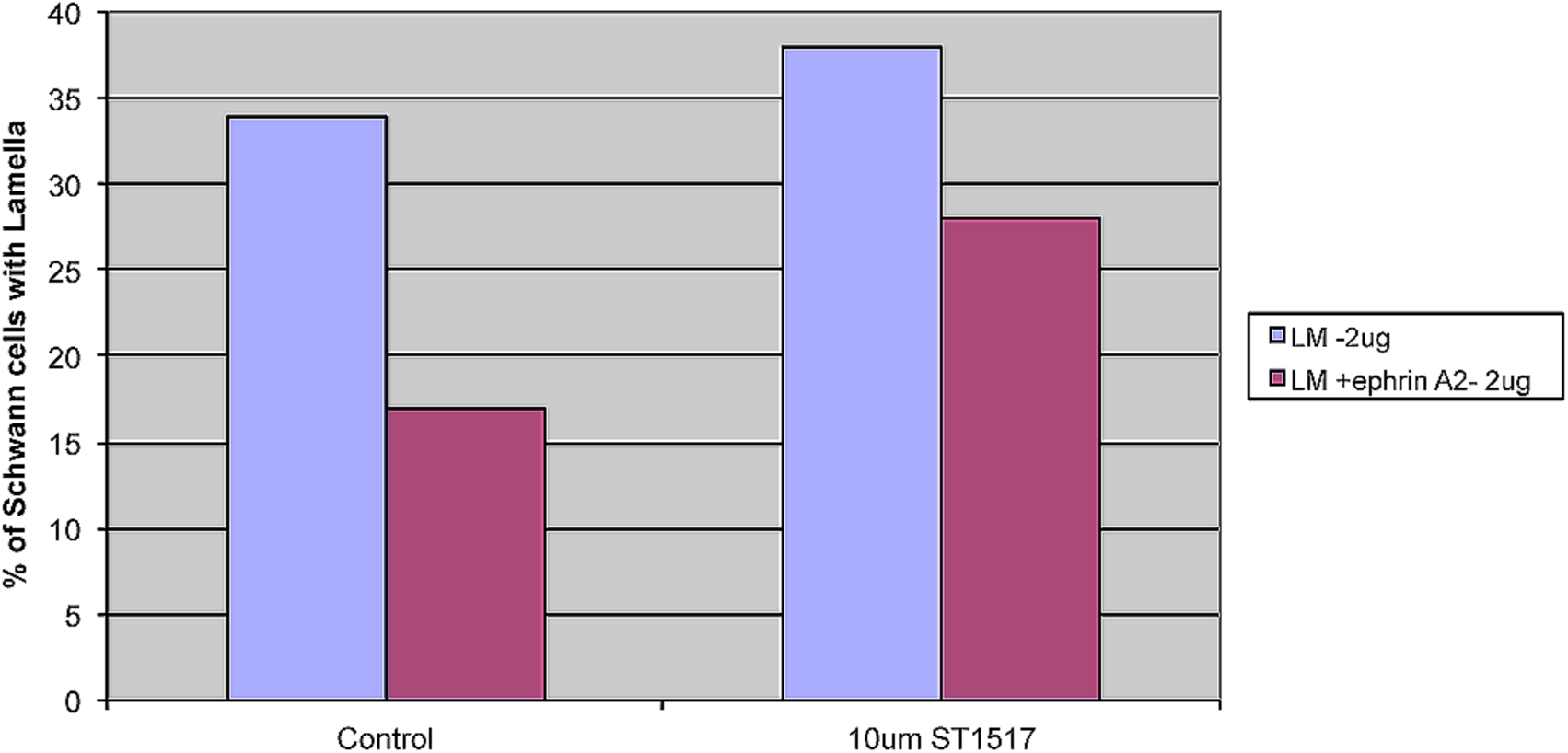
Dissociated E7 SCPs were plated on a laminin-Fc or laminin-ephrin-A2-Fc substratum for 60 minutes. Then, the percent SCPs that had flattened to form lamellar extensions was determined from a sample of microscope fields. The SCPs were plated in control medium or in medium with 10 uM of the Abelson kinase inhibitor STI571.

**Figure 14.**
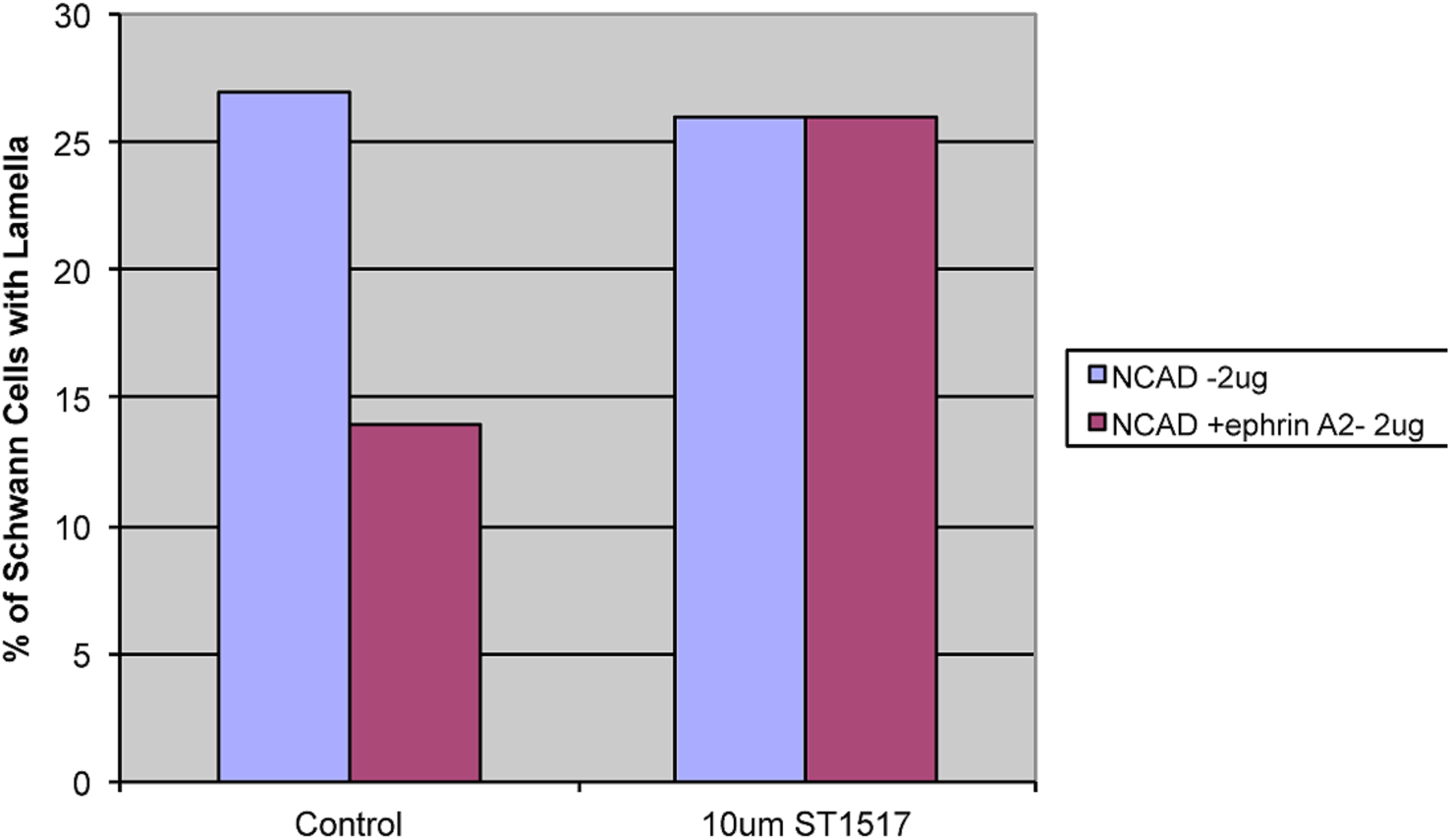
Dissociated E7 SCPs were plated on an N-cadherin-Fc or N-cadherin-ephrin-A2-Fc substratum for 60 minutes. Then, the percent SCPs that had flattened to form lamellar extensions was determined from a sample of microscope fields. The SCPs were plated in control medium or in medium with 10 uM of the Abelson kinase inhibitor STI571.

**Figure 15.**
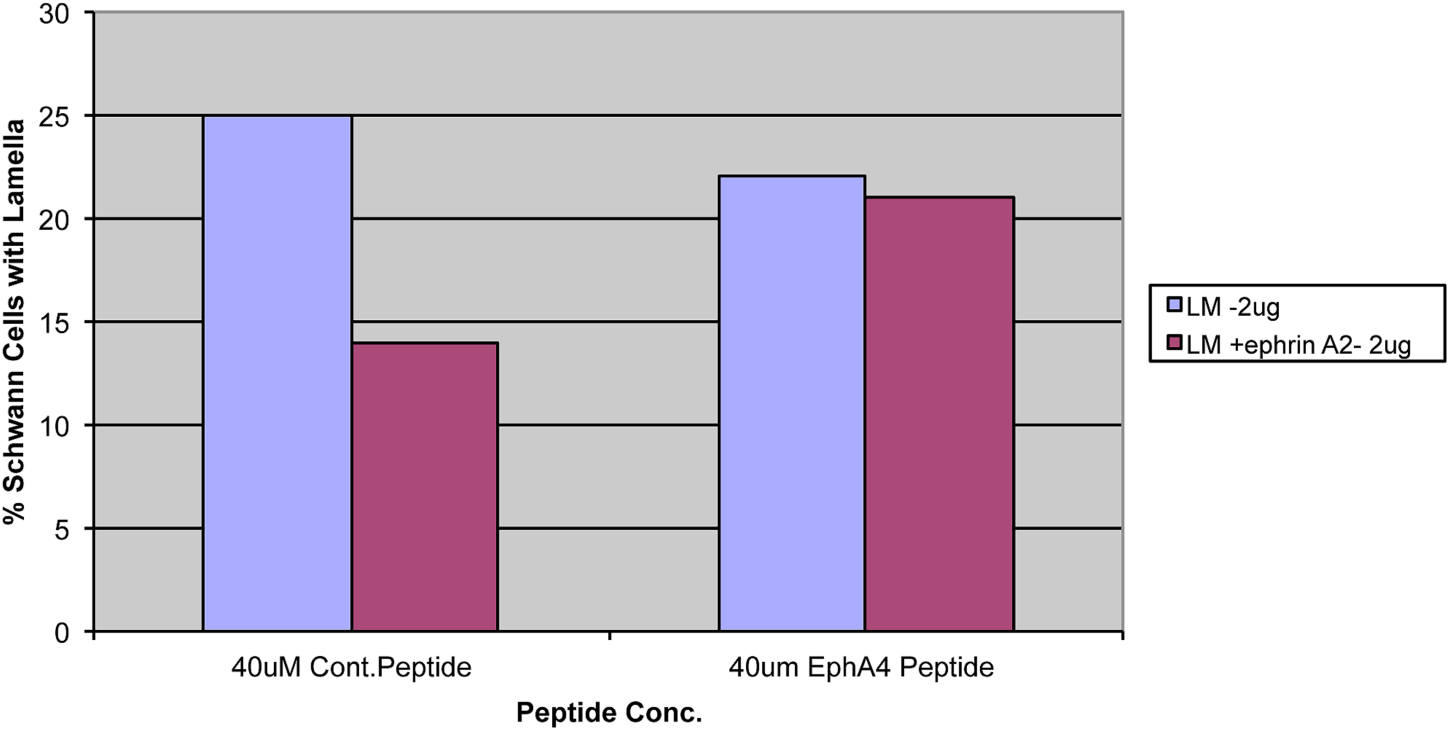
Dissociated E7 SCPs were plated on a laminin-Fc or laminin-ephrin-A2-Fc substratum for 60 minutes. Then, the percent SCPs that had flattened to form lamellar extensions was determined from a sample of microscope fields. The SCPs were plated in medium with 20 ug/ml of the KYL peptide, which blocks ephrin-A2 binding to EphA4, or with a control peptide.

**Figure 16.**
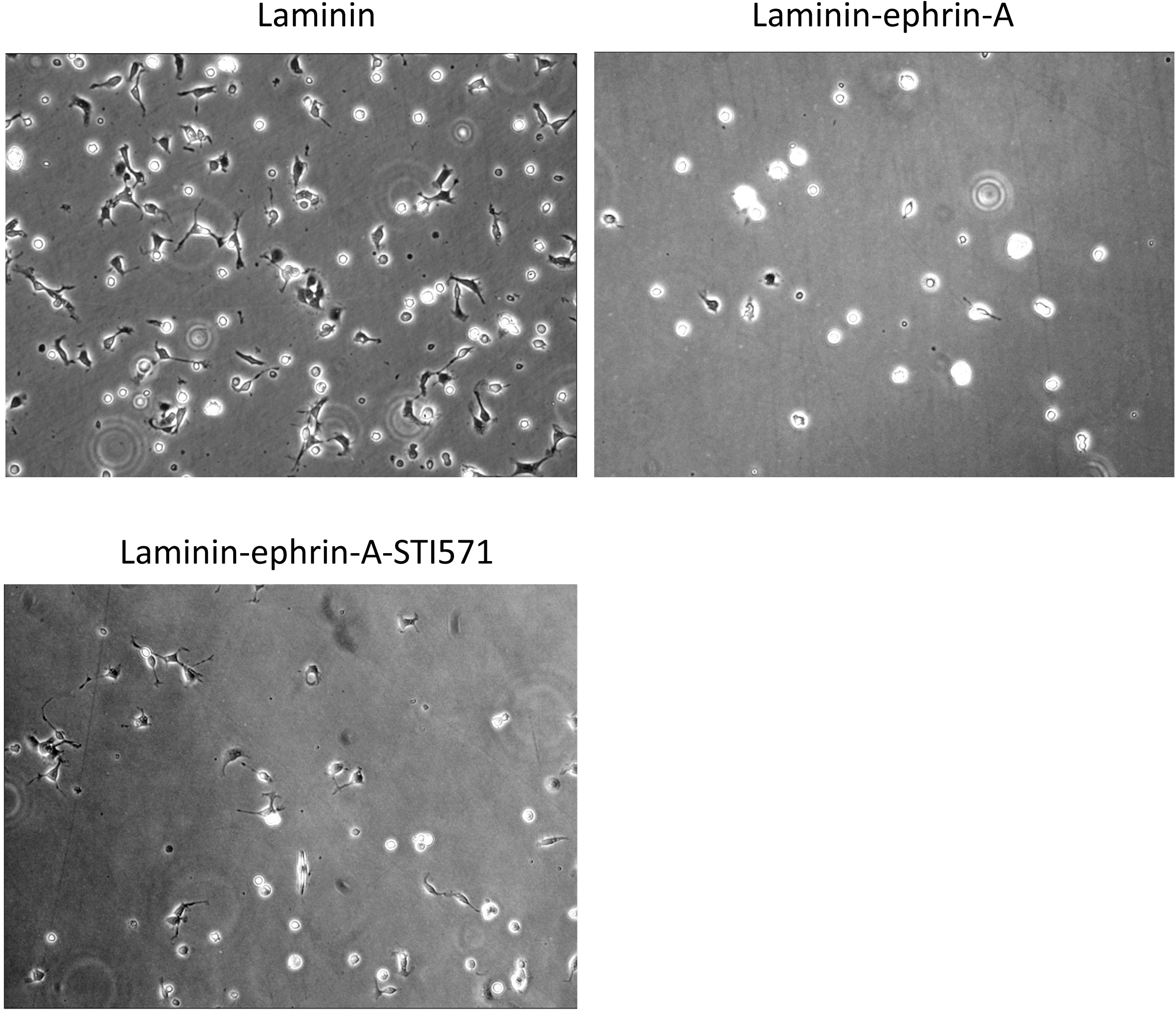
Upper panels show dissociated E7 SCPs that were plated for 60 min on a laminin-Fc or laminin-ephrin-A2-Fc substratum. Few SCPs flattened on the laminin-ephrin-A2-Fc. The lower panel shows SCPs plated on laminin-ephrin-A2-Fc with 10 uM STI571. Many more SCPs had flattened on the laminin-ephrin-A2-Fc. 20X.

As another indication that SCPs are sensitive to ephrin-A2, we transfected dissociated E7 SCPs with an ephrin-A2-GFP plasmid or a control GFP plasmid and mixed transfected cells with untransfected, dissociated E7 SCPs. We plated the cells at high density on a laminin-coated substratum. After 24 hr SCPs transfected with the control GFP-plasmid were intermixed with untransfected SCPs in a layer of mutually adherent cells (Fig. 17; left panels). On the other hand, ephrin-A2-GFP expressing cells did not join the cell layer but remained rounded or poorly intercalated among the densely packed cells (Fig. 17 right panels). These cells resembled metastatic carcinoma cells that lose contact with their primary tissue. Whether the GFP-ephrin-A2 SCP repelled their untransfected SCP neighbors by trans interactions between cells or were unable to interact because of cis ephrinA2/EphA4 interactions within transfected cells themselves, the results indicated that ephrinA2 regulates SCP adhesion and motility.

**Figure 17.**
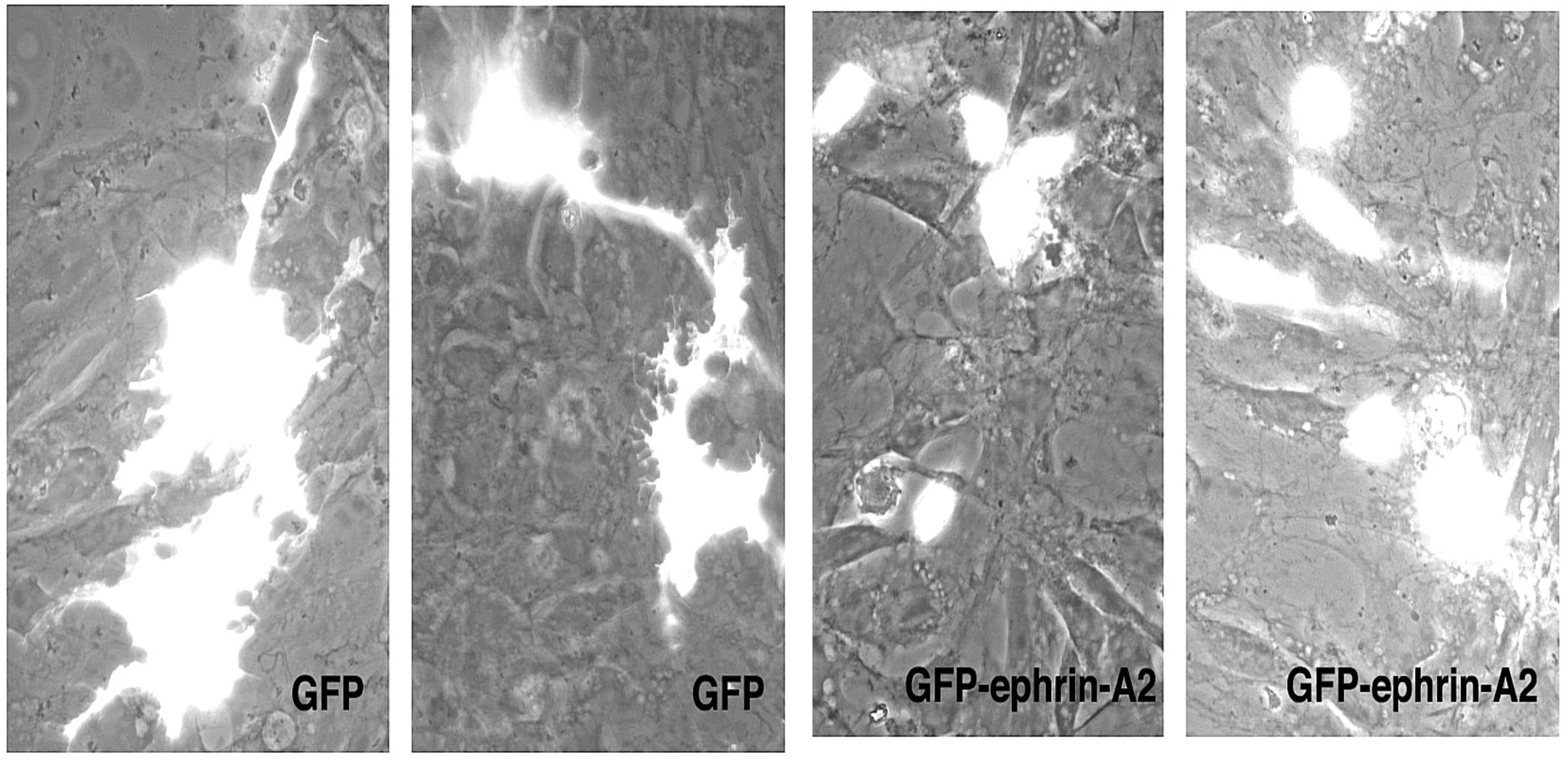
Dissociated E7 SCPs were transfected to express GFP-ephrin-A2 or a control GFP. The transfected cells were mixed with untrasfected SCPs and plated at high density on laminin. After 24 hr the control-GFP SCPs were intercalated with untransfected SCPs in a cell sheet (left panels). The ephrin-A2-GFP SCPs were excluded from the SCP cell layer or were loosely adherent to other SCPs (right panels). 40X.

Thus, these experiments indicate that ephrin-A2 signaling induces cell contractility in SCPs, and this contractility can be blocked by inhibiting ephrin-A2 binding to EphA receptors on SCPs or by inhibiting Abelson kinase activity.

### Migration of SCPs from DRG explants on laminin

When E7 DRGs were explanted onto a laminin-coated surface, hundreds of axons extended away onto the substratum. SCPs also migrated from the explants in association with the axons (Fig. 18, 19A, Video 7). Individual SCPs often adhered to and migrated along axons with a rounded cell body along an axon and an elongated cell process extending forward. Sometimes, the elongated front processes of an SCP would span between two axons. In addition, SCPs also extended flattened processes onto the laminin substratum.

**Figure 18.**
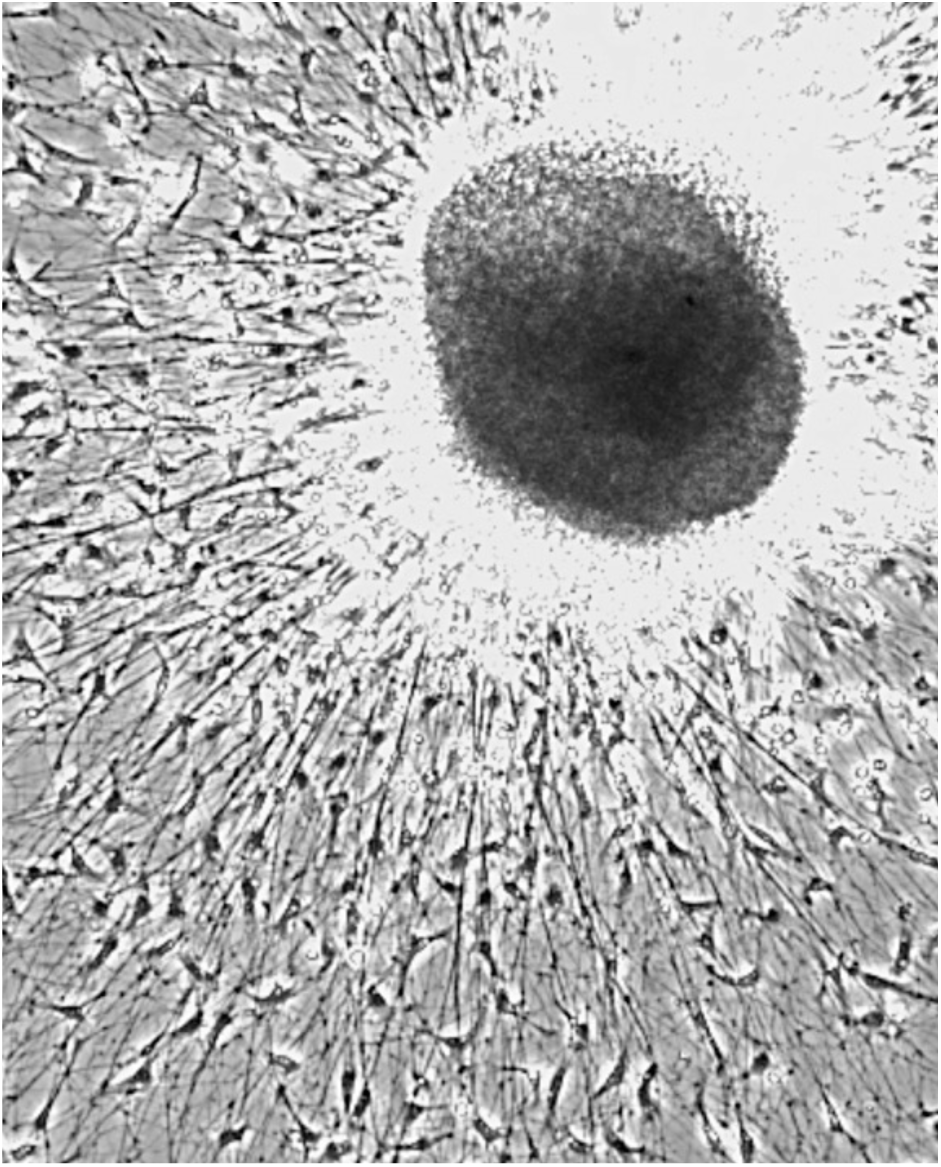
An E7 DRG explanted in high [Ca++] medium on laminin. Hundreds of SCPs have migrated out of the DRG along axons and on the laminin. 20X.

We previously showed that contact interactions between DRG neurons and SCPs depended on N-cadherin function (21,22). When DRG explants were cultured on laminin, many SCPs migrated away from DRG explants in close association with the growing axons. We determined the rates of SCP migration along axons extended from DRG explants in a variety of conditions. Time lapse videos of SCPs migrating from DRG explants were recorded for 40-60 minutes periods. The locations of randomly selected SCPs were determined and followed for 40-60 minutes at 5 minute intervals, and migration rates were determined as um/hr.

The first series of experiments DRGs examined the role of N-cadherin mediated adhesion in E7 SCP migration from E7 DRGs cultured on laminin-coated substrata. In order to remove N-cadherin function, we used media that contained a low [Ca++], when N-cadherin is not functional or media with 0.5 mM or more [Ca++], at which n-cadherin function is high. In low [Ca+] medium mean migration rate of SCPs away from DRGs was 59 um/hr, and in high [Ca++] mean SCP migration rate was 65 um/hr (Table 1).

**TABLE 1:** MIGRATION RATE OF E7 SCPs SUBSTRATUM.

|  | **SUBSTRATUM** | **CONDITION** | **RATE (um/hr)** |
| --- | --- | --- | --- |
| 1. | LAM | LO CA | 59 um/hr |
|  | LAM | HI CA | 65 um/hr |
| 2. | LAM | HI CA | 66 um/hr |
|  | LAM | HI CA + EPHA4-Fc | 28 um/hr |
| 3. | LAM | HI CA | 50 um/hr |
|  | LAM | HI CA + STI571 | 38 um/hr |
|  | LAM | LO CA | 43 um/hr |
|  | LAM | LO CA + STI571 | 57 um/hr |
| 4. | L1 | HI CA | 41 um/hr |
|  | L1 | HI CA + NRG | 57 um/hr |
| 5. | L1 | HI CA | 51 um/hr |
|  | L1 | HI CA + AG1478 | 22 um/hr |
| 6. | L1 | HI CA | 48 um/hr |
|  | L1 | HI CA + blebbistatin | 20 um/hr |
| 7. | L1 | HI CA | 38 um/hr |
|  | L1 | HI CA + H1152 | 22 um/hr |
| 8. | L1 | LO CA | 29 um/hr |
|  | L1 | HI CA | 50 um/hr |
| 9. | L1 | HI CA + control Fc | 52 um/hr |
|  | L1 | HI CA + EPHA4-Fc | 22 um/hr |
| 10. | L1 | HI CA + control peptide | 39 um/hr |
|  | L1 | HI CA + KYL peptide | 18 um/hr |
| 11. | L1 | E7 HICA | 50 um/hr |
|  | L1 | E13 HICA | 25 um/hr |
| 12. | LAM | E7 HICA | 59 um/hr |
|  | LAM | E13 HICA | 34 um/hr |
|  | LAM | E7 LOCA | 56 um/hr |
|  | LAM | E13 LOCA | 54 um/hr |

This slightly reduced rate of SCP migration in the absence of N-cadherin function suggests that sufficient adhesion to stabilize SCP extensions can involve SCP integrin receptors binding to the substratum-bound laminin and perhaps SCP binding to axons via NCAM or other axonal ligands.

We tested the role of ephrin-A2 signaling in SCP migration from DRGs by blocking ephrin-A2 binding and activation of SCP EphA4 receptors. This would inhibit the effects of axonal ephrin-A2 signaling on SCP contractility and inhibition of adhesive interactions. For these experiments we used a recombinant EphA4-Fc construct, which binds ephrin-A2 and blocks SCP EphA4-ephrin interactions. The EphA4-Fc was added when DRGs were placed in culture, and video records of SCP migration were made 24 hours later. One effect of the presence of EphA4-Fc is that fewer SCPs migrate away from the DRGs along axons. Compare Fig. 19A (control medium) with Fig. 19B (medium with 20 ug/ml EphA4-Fc). In media with high [Ca++] levels, SCP migration rate in the presence of EphA4-Fc was 41% of the control rate (Table 1, 28 um/hr vs. 66 um/hr, Video 8). We also cultured DRGs on laminin substrata in medium with the EphA4-Fc and the KYL peptide, we obtained from Dr. Pasquale (25) (Figure 20 A and B). Note in Figs. 19 and 20 how many fewer SCPs leave the DRG when ephrin-A2 binding to EphA4 is blocked, despite hundreds of axons that extended from the explant. Cell contraction seems less frequent, when ephrin-A2 signaling is blocked. These results suggest that when N-cadherin is functional, ephrin-A2 signaling from DRG axons promotes SCP migration by promoting SCP contractility and/or by reducing SCP adhesivity to N-cadherin. Without contractility it would be more difficult to release rear adhesions.

**Figure 19.**
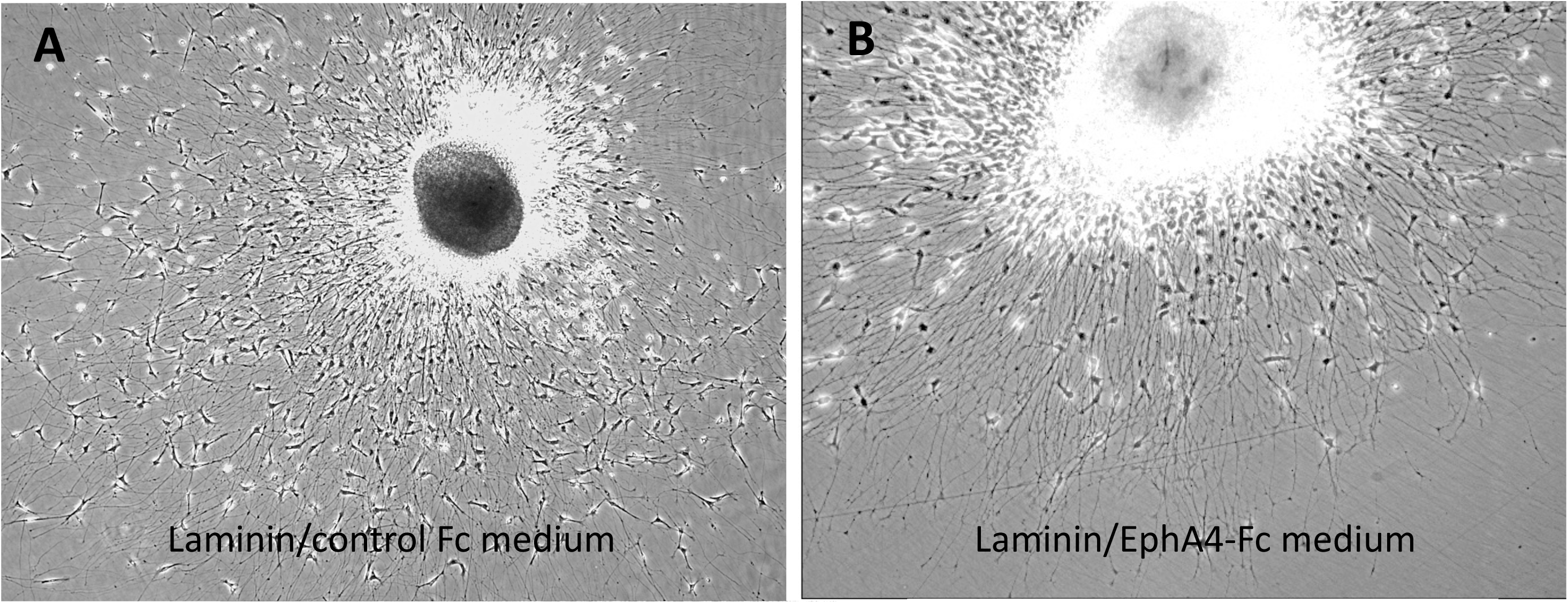
E7 DRGs cultured on laminin in high [Ca++] in (A) 20 ug/ml control-Fc medium or (B) 20 ug/ml EphA4-Fc medium. In EphA4-Fc medium fewer SCPs leave the explant. 20X.

**Figure 20.**
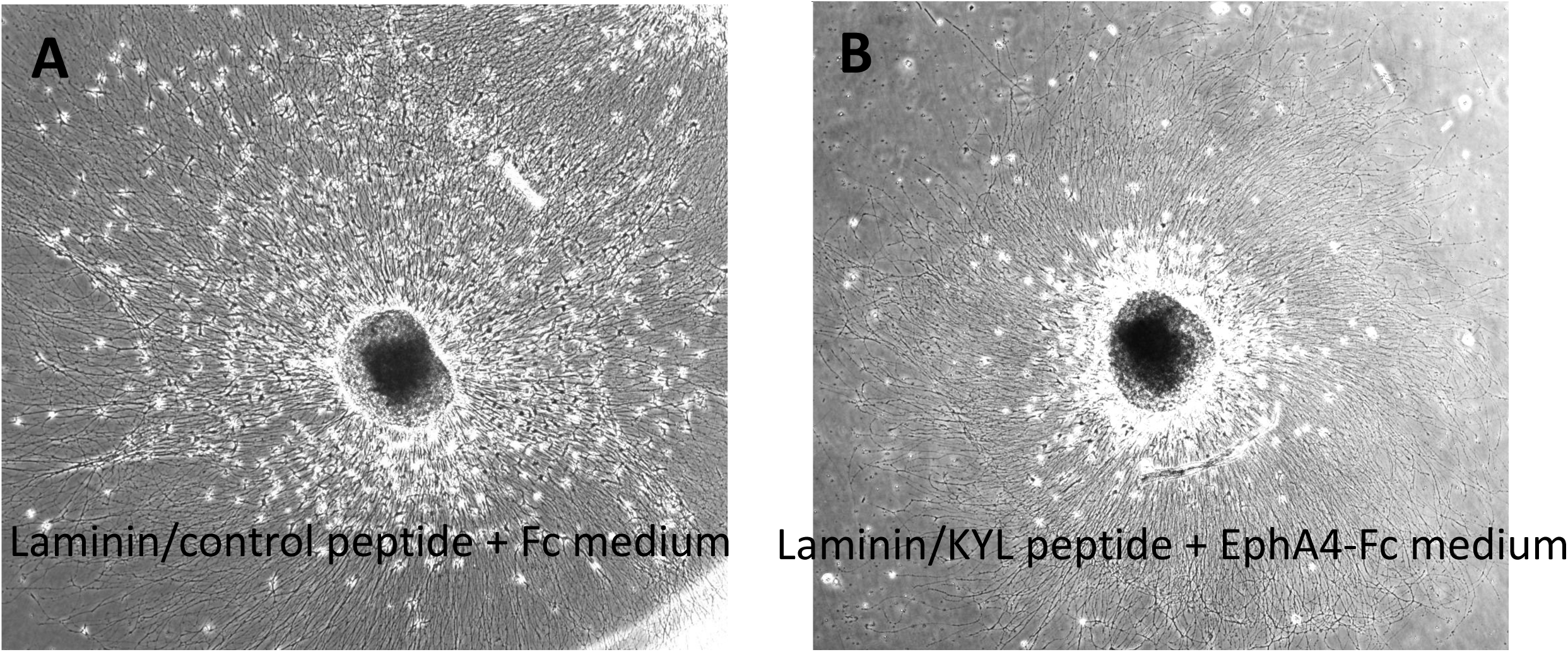
E7 DRGs cultured on laminin in high [Ca++] in (A) 20 ug/ml control-Fc and control peptide medium or (B) 20 ug/ml EphA4-Fc and 20 ug/ml KYL peptide medium. In EphA4-Fc + KYL medium fewer SCPs leave the explant. 20X.

Earlier, we increased ephrin-A2 signaling when we plated dissociated SCPs on laminin mixed with ephrin-A2-Fc. This caused reduced or slower spreading of SCPs on the substratum. Similarly, we cultured E7 DRGs on laminin-ephrin-A2-Fc substrata. Fewer SCPs migrated from DRGs onto the substratum, and SCPs that did migrate along axons were less flattened and spread on laminin-ephrin-A2-Fc (Compare Figs. 18 or 19A to Fig. 21). In the following experiments, E7 DRGs were cultured overnight on laminin substrata in low or high [Ca++] conditions (Fig. 21). Time lapse videos were recorded for 45 minutes and SCP migration rates were calculated. In high [Ca++] medium SCP migration was reduced from 55 um/hr to 38 um/hr (Table 1).. Thus, SCP migration was reduced by increased ephrin-A2 signaling.

**Figure 21.**
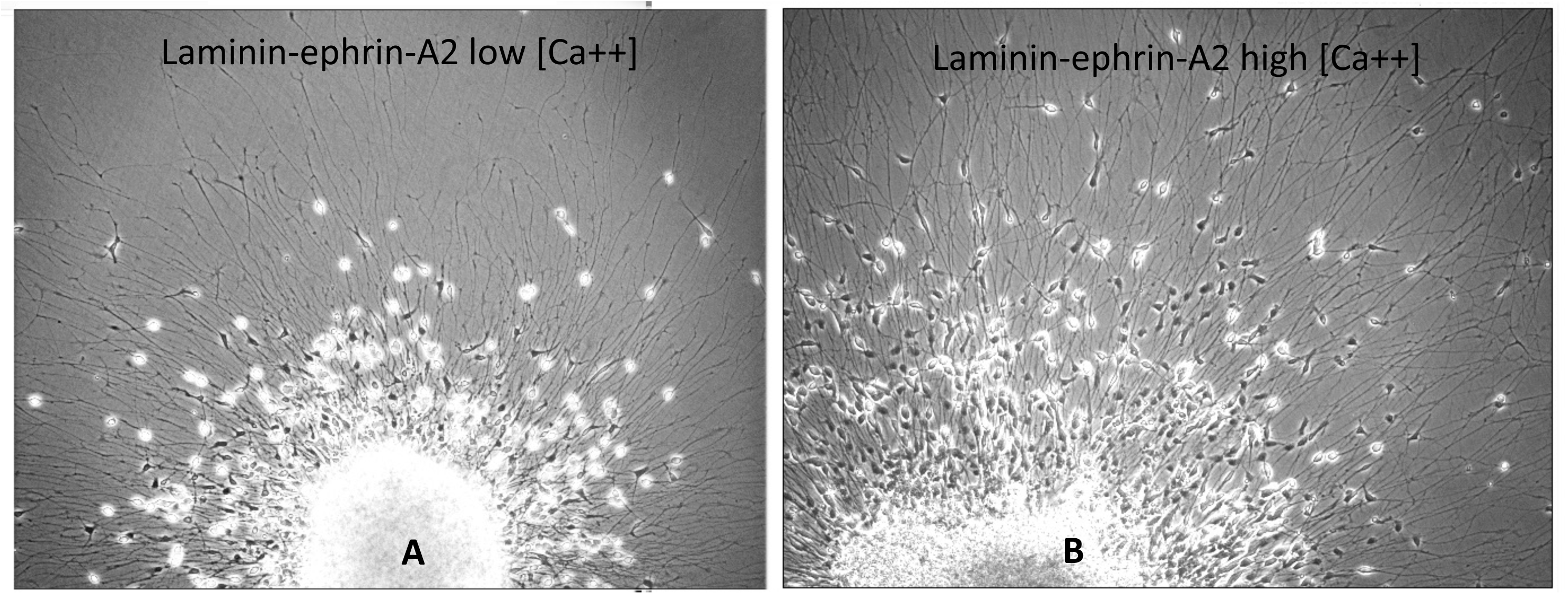
E7 DRG explanted on laminin-ephrin-A2-Fc in medium with low [Ca++], panel A, or high [Ca++] medium panel B. Fewer SCPs migrate from the DRG on laminin-ephrin-A2-Fc and many are rounded. 20X

In earlier experiments we also found that the Abelson kinase inhibitor STI571, which inhibits ephrin-A2 signaling to EphA4, blocked the reduced spreading of SCPs on a laminin-ephrin-A2 Fc substratum (Figs. 13, 16). Thus, we cultured E7 DRGs on a laminin-ephrin-A2 Fc substratum in the presence of STI571 (Fig. 22). We found that in low [Ca++], when N-cadherin function is low, SCP migration was increased from 43 um/hr to 56 um/hr by the presence of STI571 (Table 1, Video 10). Perhaps, without N-cadherin-mediated adhesion and with ephrin-A2 signaling blocked by STI571, SCP migration may be promoted by increased substratum adhesion to laminin or even by ephrin-A2/EphA4 adhesive binding between SCPs and axons that promotes SCP migration. Compare Videos 9 (N-cadherin functional) and 10 (N-cadherin not functional) and see that in the presence of STI571 SCPs are more motile on laminin in low [Ca++] medium (Video 10) compared to high [Ca++] medium (Video 9).

**Figure 22.**
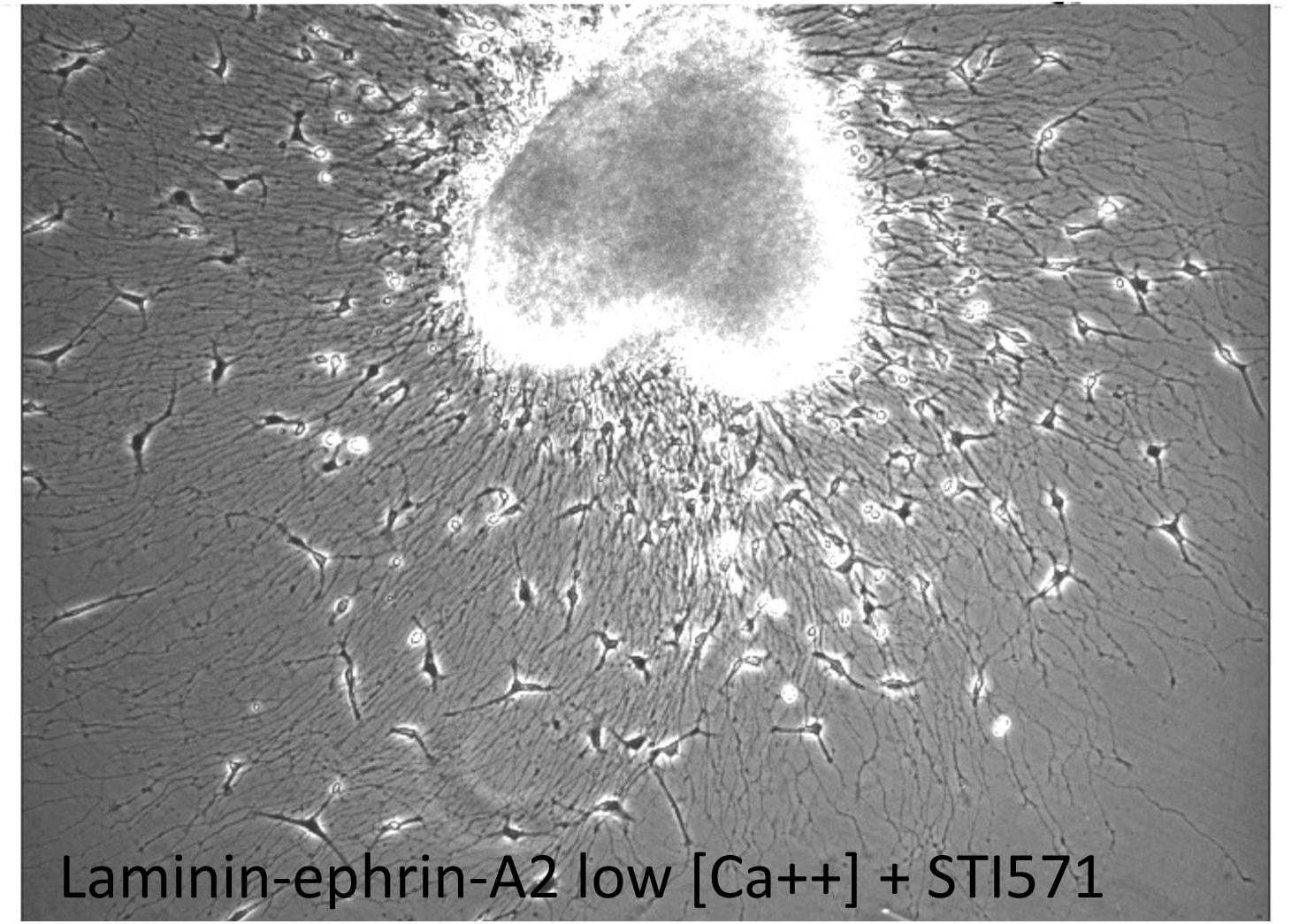
E7 DRG explanted on laminin-ephrin-A2-Fc in medium with low [Ca++] and STI571. More SCPs have left the DRG than without STI571, and they are flattened on the substrata. 20X.

### SCP migration along DRG axons on an L1- coated substratum

In the above experiments involving E7 DRGs cultured on laminin substrata, SCPs migrating away from DRG explants could adhere to and gain traction from both DRG axons and the laminin substratum. In the following experiments we cultured DRG explants on L1-coated substrata. E7 SCPs lack L1 expression (Fig. 5), and thus, cannot adhere to the L1 substratum. In this case, they gain traction for migration only from moving along axons and the surfaces of other SCPs (Fig. 23, 24, Video 11, 12, 13).

**Figure 23.**
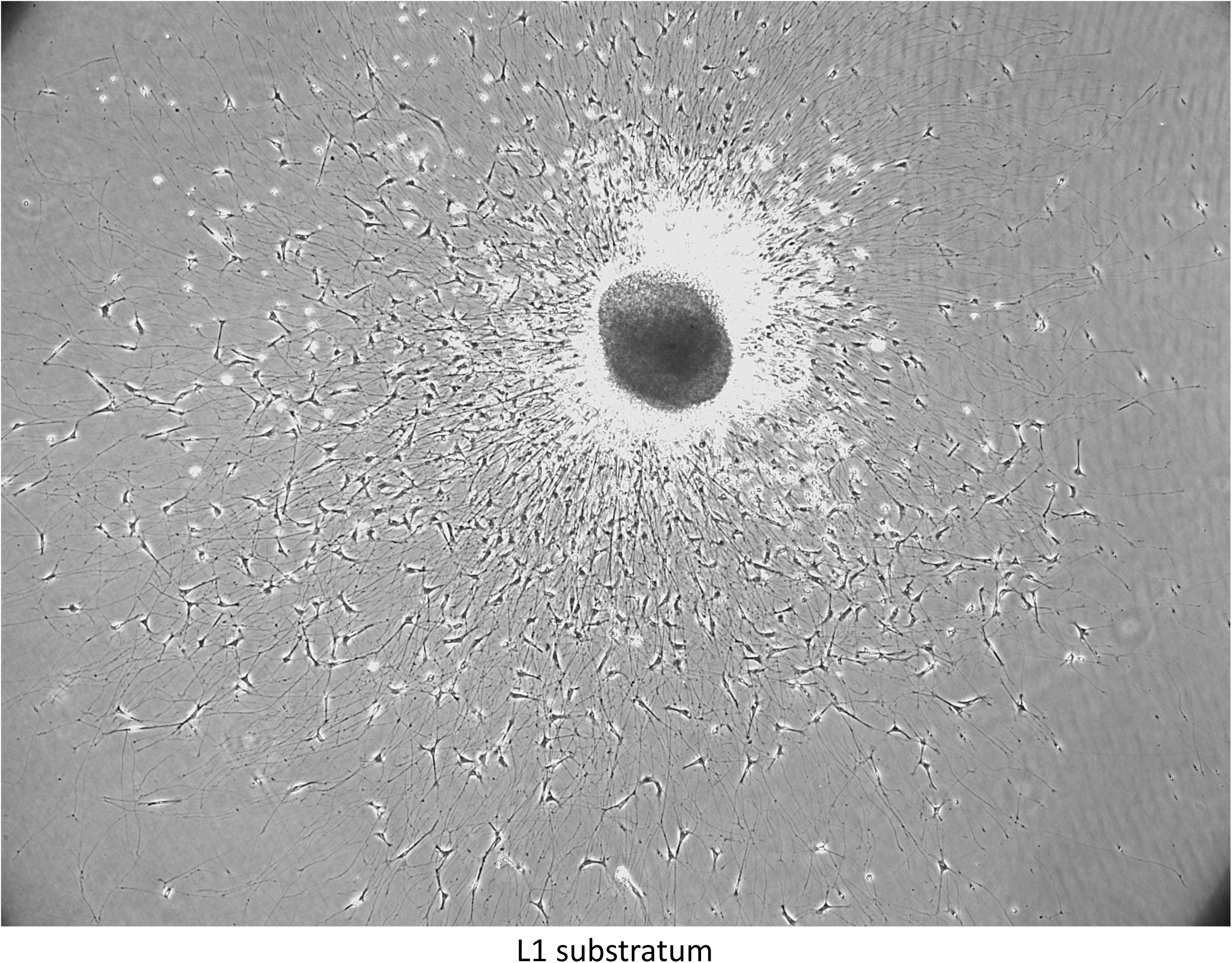
E7 DRG explanted on an L1 substratum in the presence of high Ca++ (functional N-cadherin). 20 X.

When SCPs migrate from DRGs along the growing axons, they exhibit the same dynamic anterior protrusion, adhesion and posterior contractility that drive the migration of SCP on laminin (Video 10,11). The myosin II inhibitor blebbistatin reduces the rate of SCP migration along DRG axons from 48 um/hr to 21 um/hr (Table 1). SCPs can still protrude their leading margin, but without myosin contractility net cell translocation is slower. Similarly, the SCP migration speed along DRG axons was reduced from 38 um/hr to 22 um/hr, when the Rho kinase inhibitor, H1152, was present in the culture medium (Table 1, Figure 25; Video 14). These results indicate that SCP migration along DRG axons does involve myosin-based contractility.

**Figure 24.**
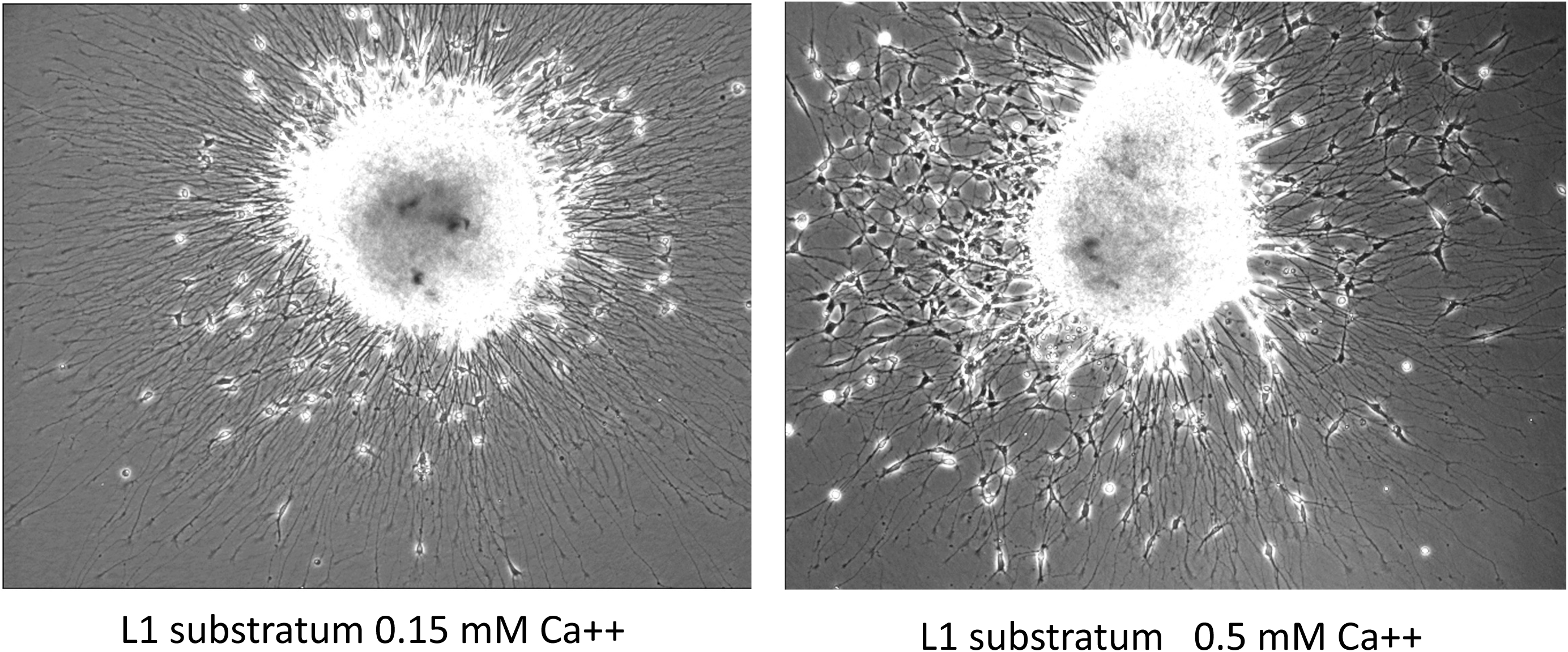
E7 DRG explanted on an L1 substratum in the presence of low [Ca++] (left panel) or high Ca++ (right panel. Many fewer SCPs migrate from the explant when N-cadherin is not functional. 20X.

**Figure 25.**
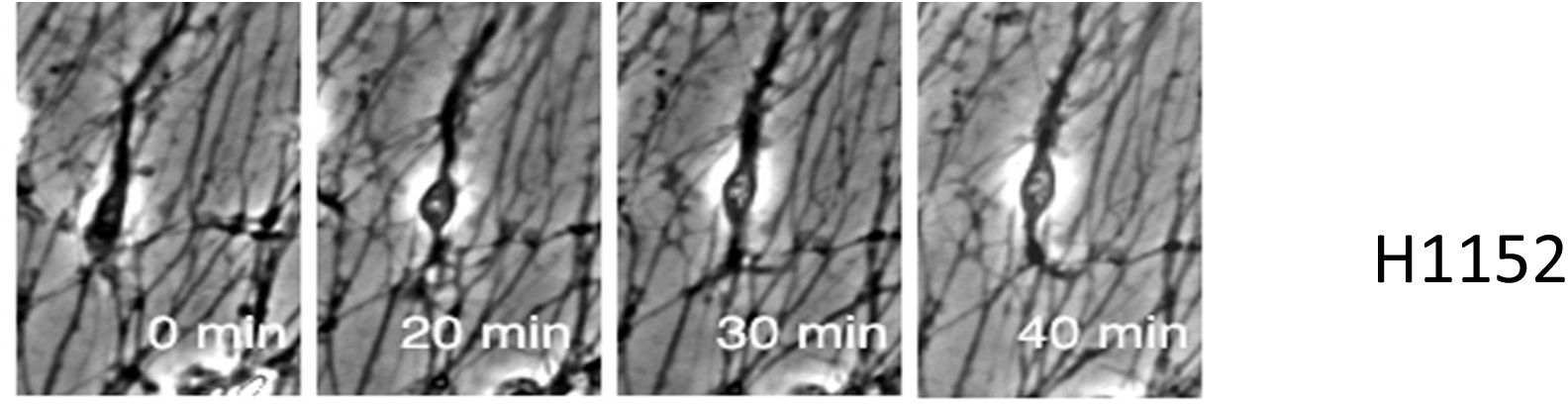
A 40 min sequence of an E7 SCP migrating on axons on an L1 substratum in the presence of the ROCK inhibitor H1152. 40X.

The expression of neuregulin1 (NRG1) type III on axons is another regulator of SCP migration along axons. We did not usually add NRG1 to the culture media of our DRG cultures, as axons express NRG1 type III on their surfaces. When we added a NRG1 protein to E7 DRG explants on L1, the rate of SCP migration along axons increased from 41 µm/hr to 57 µm/hr in the first hour after adding the NRG1 (Table 1). On the other hand, we added AG1478, an inhibitor of erbB signaling, to cultures of E7 DRG, and determined that by 30 min after adding the erbB inhibitor the rate of SCP migration along axons was reduced from 51 um/hr to 22 um/hr (Table 1). These data indicate that, like in vivo, SCP migration along axons in vitro is promoted by NRG1.

### Effects of reducing [Ca++] on SCP migration along axons on L1 substratum

We examined the effect of eliminating N-cadherin mediated adhesion on SCP migration along DRG axons, when adhesion to laminin is not available. After overnight culture of E7 DRGs on an L1-subtratum in 0.15 mM [Ca++] or 0.5 mM [Ca++], we tracked the positions of individual SCPs along axons at 5 minute intervals for 60 minutes. Many fewer SCPs migrated along axons extended from the DRG explant in 0.15 mM [Ca++] Fig. 24. When [Ca++] was 0.5 mM or more, N-cadherin was functional and the mean rate of SCP migration along axons was 50 µm/hr (Fig.26; Video 12). When the DRGs were cultured in a low [Ca++] medium (0.15 mM), when N-cadherin is non-functional, SCP migration along axons on an L1 substratum was reduced to 29 µm/hr (Table 1, Fig. 27, Video 13). This result shows that N-cadherin-mediated adhesion of SCPs to axons is important in SCP migration along E7 axons.

**Figure 26.**
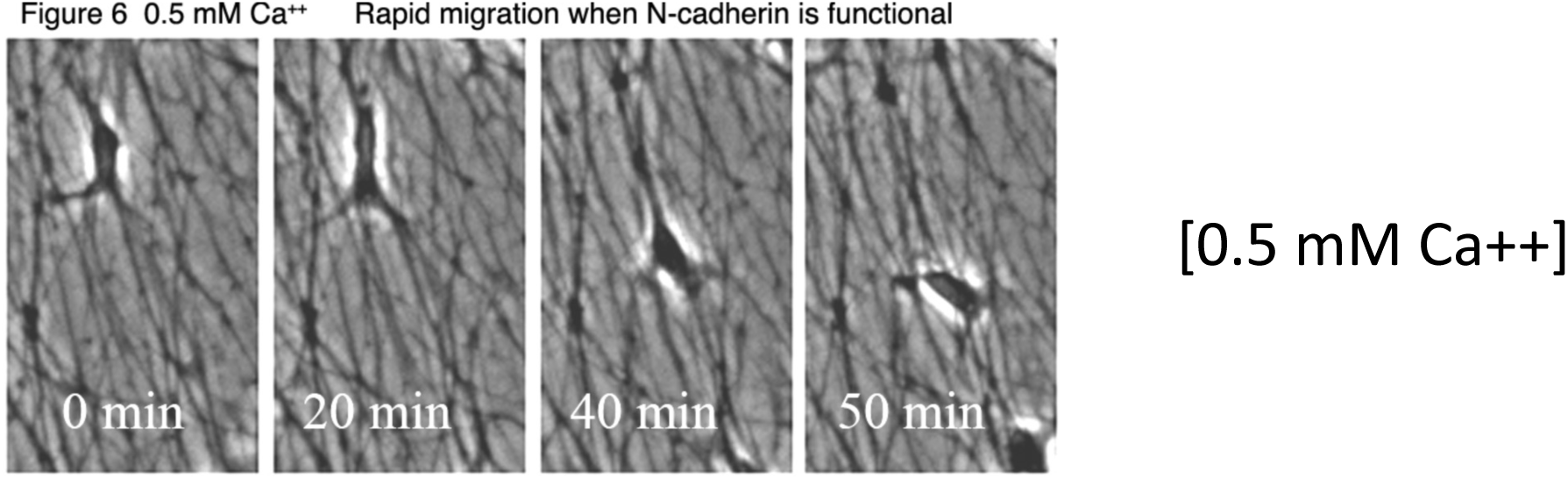
A 50 min sequence of an E7 SCP migrating on axons on an L1 substratum in high [Ca++] medium with functional N-cadherin. 40X.

**Figure 27.**
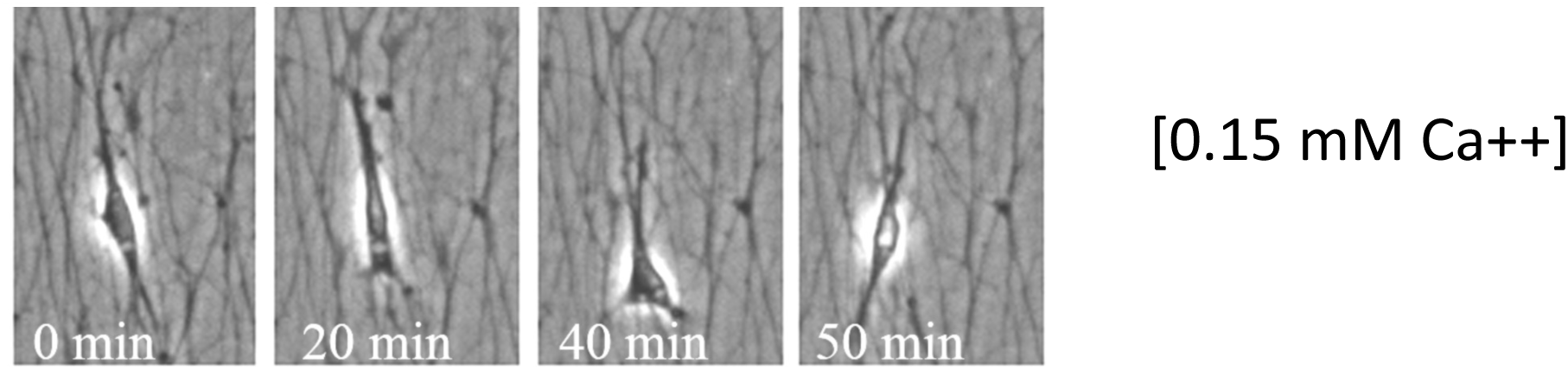
A 50 min sequence of an E7 SCP migrating on axons on an L1 substratum in low [Ca++] medium without functional N-cadherin. 40X.

### Ephrin-A/EphA interactions regulate SCP migration along axons

The above experiments showed that SCP migration is reduced without N-cadherin function. We next conducted experiments to remove signaling from axonal ephrin-A2 to EphA4 receptors on SCPs. Our hypothesis predicts that blocking ephrin-A2-EphA4 interactions will reduce contractility and/or increase SCP-axon adhesion, resulting in reduced SCP migration rate.

As we did on laminin substrata, we inhibited EphA signaling by adding EphA-Fc chimeric proteins that competitively block axonal ephrin-A ligands from binding to SCP EphA receptors. We added 10 µg/ml EphA3-Fc or control Fc to DRG explants and at 24 hr we measured SCP migration along axons. The rate of SCP migration along axons in the presence of control Fc was 52 µm/hr, while the rate was slowed to 22 µm/hr in the presence of EphA3-Fc (Table 1, Fig. 28; Video 15). This is consistent with our hypothesis and implicates EphAs and ephrin-As in regulating SCP migration along axons. However, these data do not implicate specific EphAs or ephrin-As. In the following experiments we used a selective approach to block activation of EphA4.

**Figure 28.**
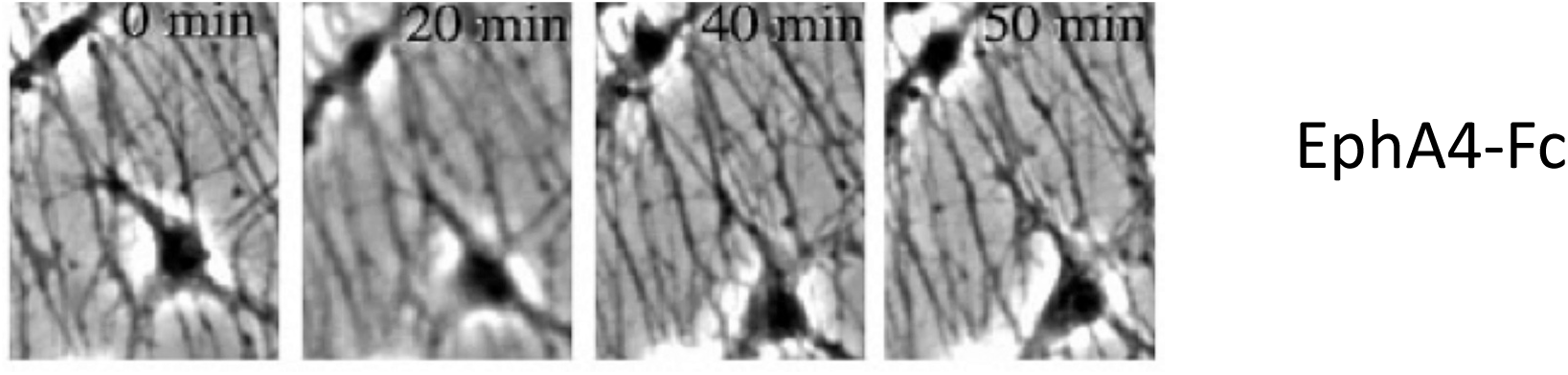
A 50 min sequence of E7 SCPs migrating on axons on an L1 substratum in high [Ca++] in the presence of 20 ug/ml EphA4-Fc, which blocks ephrin-A2 signaling. 40X.

We used the KYL peptide obtained from Dr. Pasquale that binds tightly and selectivity to the ephrin-A binding site of EphA4. E7 DRG explants were cultured on L1 for 24 hr, and time lapse images were recorded 60 min before and at 2 hr after adding the EphA-blocking peptide or the control peptide. After adding the EphA4-blocking peptide, the rate of SCP migration along axons decreased from 39 µm/hr to 18 µm/hr (Table 1, while the control peptide did not inhibit SCP migration. In another experiment E7 DRGs were cultured 24hr in the presence of the KYL peptide or control peptide, and then time lapse images were recorded for 60 min. Few SCP had migrated from the explants treated with the KYL peptide. To analyze migration, we drew a 45 µm circle (about one cell diameter) around individual SCP and determined whether the cell migrated out of the circle over the 60 min period. In the presence of the control peptide 76% of SCP migrated out of the circle along axons, while in the presence of the EphA4-blocking peptide only 33% of SCP migrated out of the 45 µm circle over 60 min.

### SCP migrate along E13 axons slower than along E7 axons

Our immunocytochemical studies indicated that as DRG neurons mature from E7 to E13, axonal expression of NCAM expression decreased, and N-cadherin expression increased on E13 axons. L1 was expressed on axons at both times (Fig. 3). In addition, expression of ephrin-A2 decreased substantially on E13 axons (Fig. 4). Our model suggests that because of reduced ephrin-A2-EphA4 signaling, and increased N-cadherin-mediated adhesion to E13 axons, E7 SCPs may migrate slower along E13 DRG axons, compared to E7 axons. To test this, we prepared SCPs from E7 peripheral nerves, marked them with a fluorescent cell tracker, and added fluorescent E7 SCPs to E7 and E13 DRG explants on an L1 substratum (Video 16). After 24 hr we located fluorescent E7 SCPs and recorded their migration along E7 axons or E13 axons (Fig. 29, 31 A and B are at beginning of video 17, C and D are at end of Video 17). We found that the mean migration rate of fluorescent E7 SCP on E7 axons was 50 µm/hr, while the migration rate of E7 SCP along E13 axons was 25 µm/hr (Table 1). These results indicate that with increased N-cadherin expression on E13 axons, adhesion of E7 SCPs to E13 axons is increased over adhesion to E7 axons, and because of reduced contractility from absence of ephrin-A2, or because posterior adhesions can’t be easily released without ephrin-A2 signaling, migration along E13 axons is slowed.

**Figure 29.**
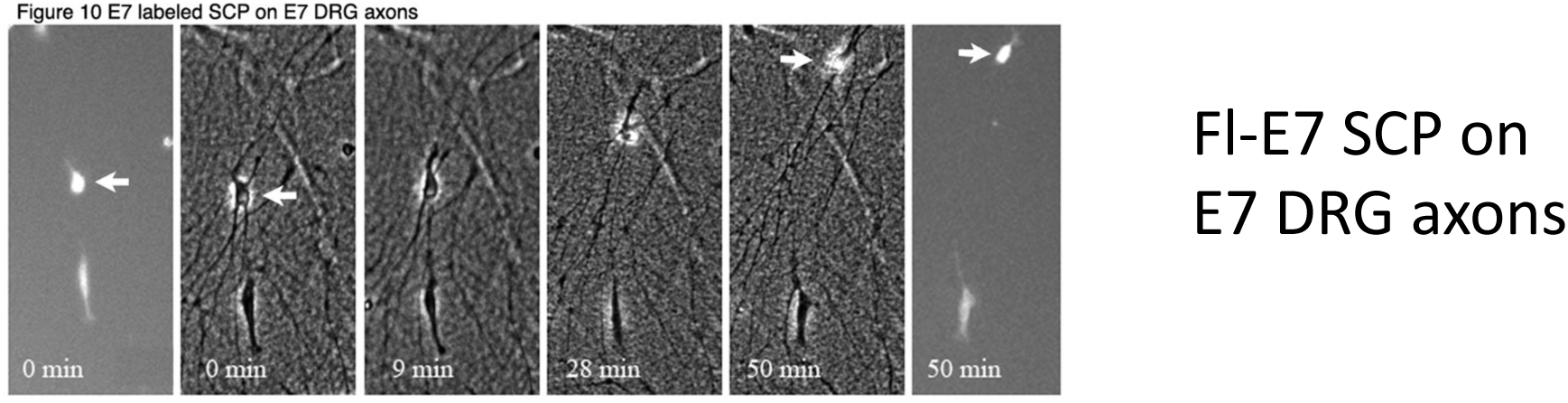
A 50 min sequence of a fluorescently labeled E7 SCP migrating on unlabeled axons of an E7 DRG on an L1 substratum in the presence of high [Ca++]. 40X.

We also cultured fluorescently marked E7 SCPs with E7 or E13 DRGs on laminin-coated substrata. We found that similar to our results on an L1 substratum, in medium with functional N-cadherin, E7 SCPs migration along E13 axons was significantly slower than migration of fluorescent E7 SCPs along E7 DRG axons (Table 1, Fig. 31, Video 18). However, at low [Ca++] when N-cadherin function is absent, the migration rates of E7 SCPs along E7 and E13 axons were the same and were faster than E7 SCP migration on E13 axons, at a [Ca++] when N-cadherin is functional (Table 1). These results also suggest that in the absence of ephrin-A2 signaling on E13 axons than the slower migration of E7 SCPs along E13 DRG axons was due to high N-cadherin-mediated adhesion of E7 SCPs to E13 axons, compared to E7 axons.

**Figure 30.**
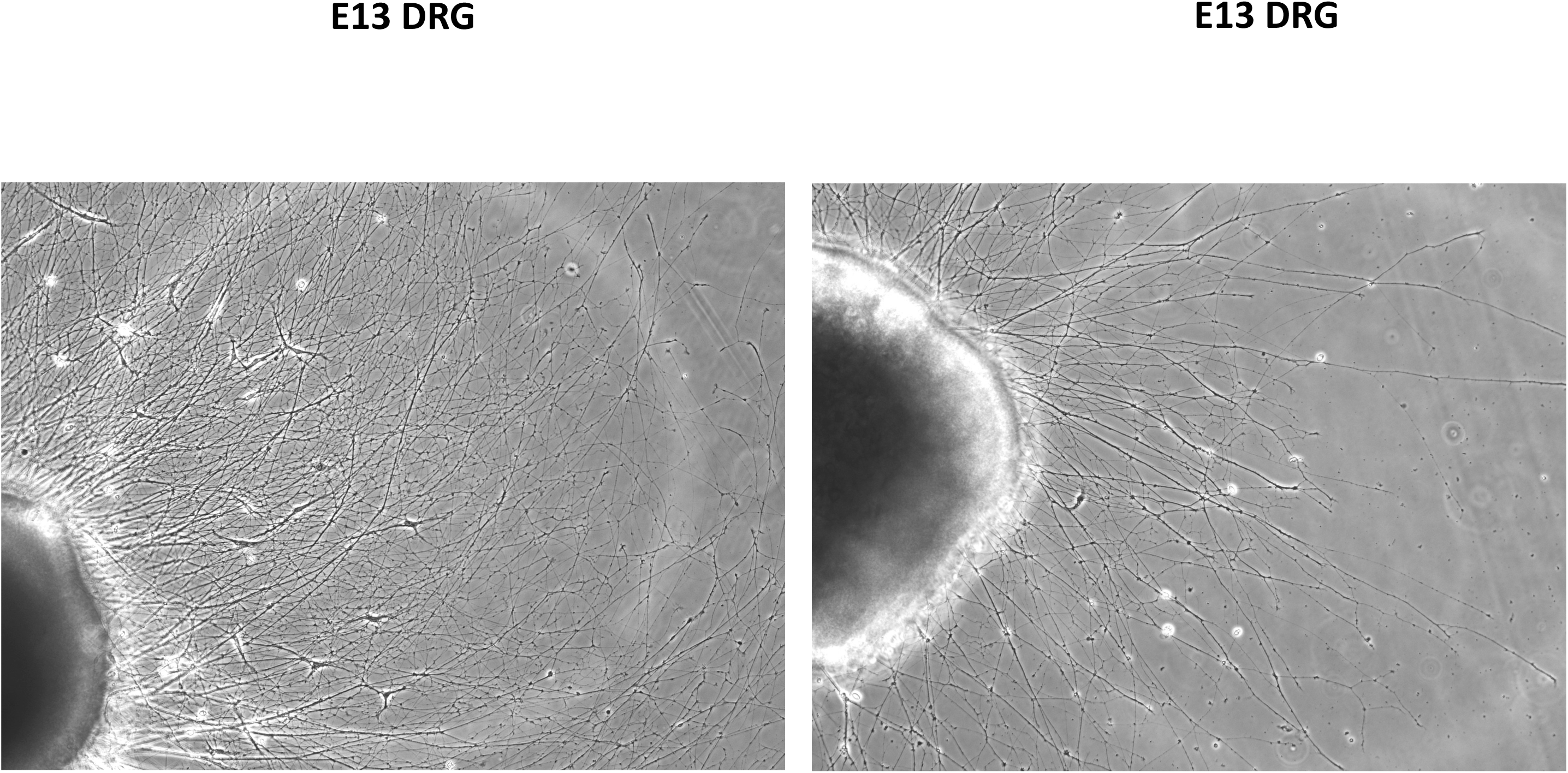
Phase contrast images of E13 DRGs cultured on laminin for 24 hr. Many fewer cells have migrated from the explants along with the axons, compared to E7 DRGs. Compare with Figures 18, 19A, 20A. Kindly provided by Dr. Andrea Ketchsek, Assistant Professor at Arcadia University,

**Figure 31.**
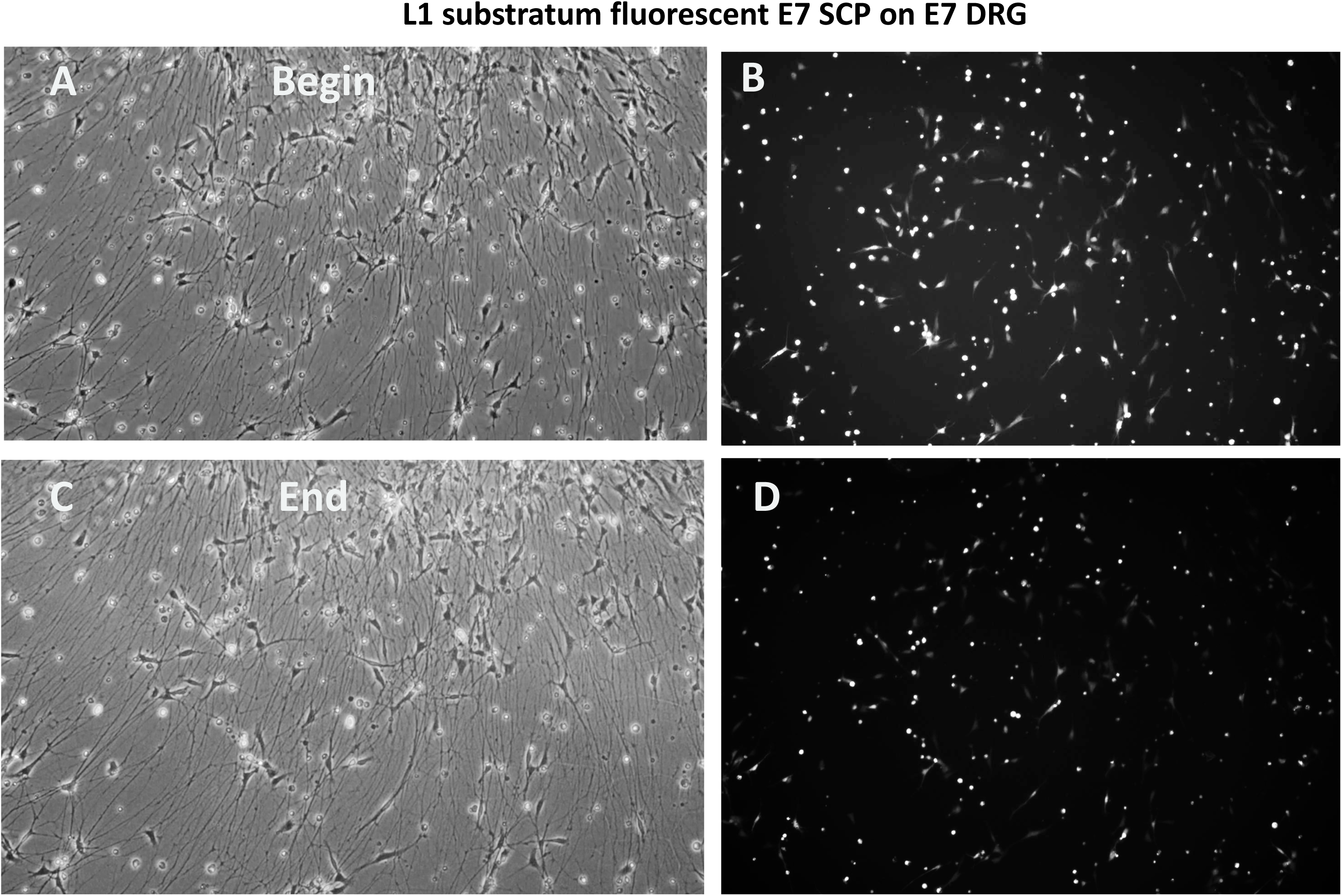
Phase contrast and fluorescence images at the beginning and end of Video 17, which shows fluorescently labeled E7 SCPs migrating on unlabeled axons of an E7 DRG on an L1 substratum in high [Ca++] medium. 20X.

## Discussion

In these preliminary studies we investigated the roles of N-cadherin-mediated cell adhesion and ephrin-A2/EphA4-mediated cell contractility in regulating the migration of E7 Schwann cell precursors (SCPs) along E7 DRG axons. At E7 chick limbs are rapidly growing and lengthening, and in the limbs’ peripheral sensory nerves, SCPs are migrating and axons are elongating from DRGs to peripheral targets (3,39,40). Our results indicate SCP migration during this phase of rapid limb growth involves a balance of adhesion and contraction to promote SCP migration along axons, as described in the cell migration model, presented in the Introduction (6,20,26).

The associations between SCPs and growing axons during the initial growth of peripheral nerves are well documented (3,39,40). During this period, SCPs are close and consistent companions to growing axons all the way to the nerve front. SCPs are the main substratum for growth cone migration. N-cadherin plays a key role in establishing and maintaining these relationships during development. SCPs may extend slightly ahead of growth cones, but their dependence on axon-derived neuregulin for survival restricts SCPs from moving ahead of growth cones. SCPs do not survive in axonless limbs (3).

Ephrins are often inhibitory repulsive guidance cues in regulating behaviors of neural crest cells during development, and in regulating Schwann cell behaviors after nerve injury (4,10,41). The repulsive effects of localized ephrins direct the early migration of neural crest cells (9,27,29). After spinal cord injury astrocytes produce ephrin-A5 that channels Schwann cell movements (42). After injury, ephrin-B2 produced by nerve fibroblasts restricts Schwann cell movements, so they cluster into groups that provide a bridge for axonal regeneration . After CNS injury ephrin-B3 produced by myelin stops Schwann cells from mingling with myelin, directing Schwann cells to migrate along blood vessels into injury sites. However, in an in vitro study ephrin-A2 promoted Schwann cell migration on a matrigel substratum, perhaps by promoting cell contractility.

We investigated the role of N-cadherin-mediated adhesion in E7 SCP migration along E7 axons by comparing the rate of SCP migration at a [Ca++] that eliminated N-cadherin function vs. a normal [Ca++] at which N-cadherin adhesive function was active. We found that when N-cadherin function was absent, E7 SCP migration rate along axons was reduced. This was more true on an L1 substratum, where E7 SCPs do not adhere to the substratum, and much less so on laminin, where in addition to N-cadherin on axons, SCP can adhere to laminin for cell spreading and traction. These results suggest that the first step in cell migration, advance of the leading cell margin of SCPs along axons, is promoted by N-cadherin-mediated binding of the SCP leading edge to axons.

As a counterpart to the investigation of N-cadherin mediated adhesion in SCP migration, we investigated the role of ephrin-A2 signaling in SCP migration along axons. Our experiments with E7 SCPs from peripheral nerve segments showed that localized ephrin-A2 signaling induced local cell contraction, and substratum-bound ephrin-A2 inhibited SCP spreading. Because E7 DRG axons express ephrin-A2, we blocked ephrin-A2 signaling to EphA4 receptors on SCPs and assessed the effects on SCP migration on E7 axons. We found that E7 SCP migration along E7 axons was inhibited when we blocked ephrin-A2 signaling to SCPs in three ways (EphA4-Fc, KYL peptide, and STI571. This result indicates that ephrin-A2 signaling promotes E7 SCP migration along E7 axons.

E13 axons express higher levels of N-cadherin and lower levels of ephrin-A2 than E7 axons, so we compared migration rates of E7 SCPs on E7 vs. E13 axons. On both L1 and laminin substrata at normal [Ca++] the migration of E7 SCPs was slower on E13 axons than on E7 axons. However, also on a laminin substratum, we found that the E7 SCP migration rate on E13 axons was faster when [Ca++] was low, which eliminated N-cadherin function. This result suggests that adhesion of E7 SCPs to E13 axons was so high at normal [Ca++] that SCP migration was inhibited by too much adhesion. Similarly, when we blocked ephrin-A2 signaling with STI571, we found that E7 SCPs migrated faster from E7 DRGs on laminin in low [Ca++] medium, when N-cadherin was not functional. These results suggest that a high level of N-cadherin mediated adhesion slows the third phase of cell migration, the release of posterior adhesions.

## Summary

As presented in the introduction to this study, optimal cell migration requires first adhesion to stabilize forward extensions of a cell, then contraction to pull the cell forward and lastly the release of posterior adhesions. We investigated the roles of both n-cadherin mediated adhesion and ephrin-A2-induced contractility in regulating the migration of E7 SCPs along DRG axons. We found that SCP adhesion to axons via N-cadherin provides substratum adhesion that promotes SCP migration, as loss of N-cadherin adhesion reduced SCP migration along axons on an L1 substratum. However, too much N-cadherin adhesion (E7 SCPs on E13 axons) may inhibit SCP migration. We showed that ephrin-A2 signals cause SCP contractility and that blocking ephrin-A2 signaling from axons to SCPs inhibited SCP migration along axons. In the Introduction to this paper we mentioned a model for cell migration that proposes that speed of cell migration depends in a biphasic manner on cell-substratum attachment strength with maximum migration at intermediate levels of cell-substratum adhesiveness, which allows formation of new adhesions at the cell front, but also allows rupture of posterior attachments powered by cell contractility. In this study we found that when N-cadherin adhesion is absent, SCP migration is slower (low [Ca++] on an L1 substratum, Fig. 28, Table 1) but when N-cadherin-mediated adhesion is too high, SCP migration is also reduced (compare E7 SCPs on E13 axons at high vs. low [Ca++], Table 1, Fig. 32, Video 15). Likewise, we found that when ephrin-A2 signaling is blocked, SCP migration is reduced (Table 1, Figs. 18, 19, 29, Video 7, 13), but when ephrin-A2 signaling is increased, SCP migration is also reduced (Table 1, Figs. 21, 22, 23, Video 8). Thus, at E7 SCP migration involves a balance of N-cadherin and integrin-mediated adhesive interactions with axons and extracellular matrix and ephrin-A2 signaling that induces cell contractility and maybe reduces adhesivity.

**Figure 32.**
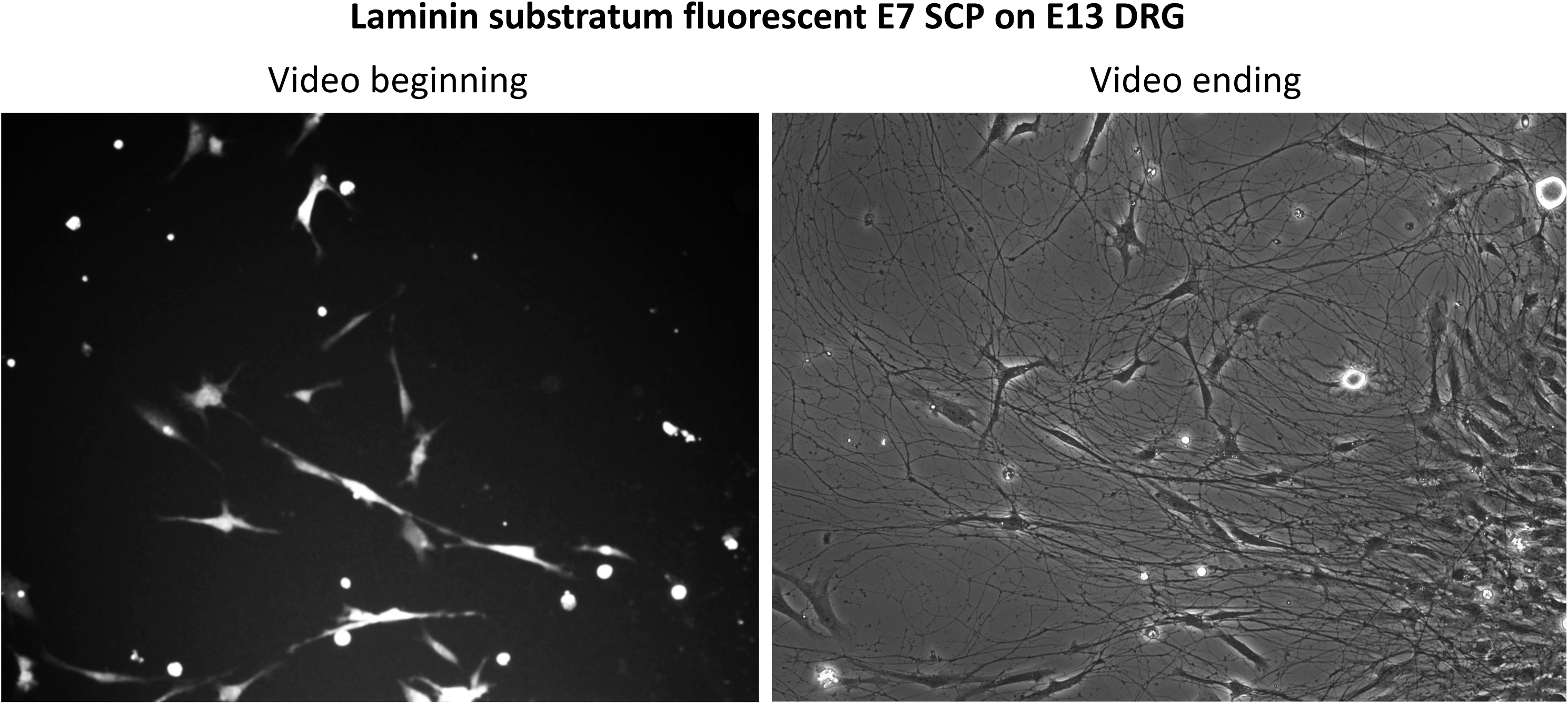
Fluorescence image (left panel) taken at the beginning of Video 18, and phase contrast image taken at the end of Video 18 of fluorescent E7 SCPs migrating on an E13 DRG on a laminin substratum in high [Ca++] medium. 20X.

## Methods

### Materials

F-12 medium, DMEM medium, Calcium-free DMEM, B27 additives, laminin, fibronectin, poly-D-lysine, Alexa Fluor 488–phalloidin and Alexa Flour 568 secondary antibodies, and CellTracker dye were purchased from ThermoFisher Scientific. NGF, neuregulin, L1–Fc, ephrin-A2–Fc, EphA4-Fc, and Fc recombinant proteins were purchased from R&D Systems. Antibodies to localize myosin II, HNK1, 1E8, p75, L1, NCAM, N-cadherin, ephrin-A2 and -A5 and EphA4 were purchased from R & D Systems and Santa Cruz Biotechnology Inc. H1152, blebbistatin, AG1478 and STI571 were purchased from Biomol.

### Preparation of DRG and peripheral nerve cultures

Dorsal root ganglia (DRG) were dissected from E7 and E13 chick embryos. Peripheral nerve segments, distal to the DRGs were also dissected from E7 and E13 chick embryos. To obtain SCPs, peripheral nerve segments were trypsin-dissociated in the presence of Ca^++^ to retain surface N-cadherin. To deplete the cultures of fibroblasts, the cell suspensions were pre-plated on fibronectin-treated (50 ug/ml for 4 hr) tissue culture dishes for 1 hr, and the supernatant with unattached cells was removed, pelleted, and plated in the experimental dishes. For cell tracking experiments SCPs were incubated with CellTracker dye according to instructions.

### Immunocytochemistry

DRGs, peripheral nerve explants and dissociated SCPs were cultured on laminin-coated coverslips in F12 medium, with added B27 supplement, and 10 ng/ml nerve growth factor (NGF; R & D Systems) for the DRGs and 10 ng/ml recombinant neuregulin (R & D Systems) for the peripheral nerve explants. After 24 hr culture, the explants and cells were fixed with 4% paraformaldehyde, 5 % sucrose in PBS, pH 7.4 for 30 min at 37° C. The coverslips were processed for immunocytochemistry, according to methods we routinely used.

### Image Analysis

The levels of expression of cell surface proteins, ephrins, and Ephs were determined from the fluorescence intensity measured in equivalent areas of stained axons and SCP, using the Metamorph program (Universal Imaging Corp; 78). For sampling along axons the line tool was used to define a 20 µm line along which was computed the mean fluorescence intensity per pixel. With the computer mouse the same line was moved slightly outside the cell, where the mean fluorescence of the line over the background was measured and subtracted from the fluorescence signal of the stained axon. For SCPs and immature SCs the Metamorph tool was used to define a 3 µm box over the flat lamellar portion of a random sample of cells, where the mean fluorescence intensity per pixel was determined. The small box was moved outside the cell boundary to determine background fluorescence, which will be subtracted from the fluorescence intensity measured on the SCP lamella. The Excel program was used to determine totals and averages.

### SCP migration assays

Microscope fields of axons with SCPs were chosen using a 20X or 40X objective on an Olympus inverted microscope. Images of a microscope field were captured every 60 sec-5 min for 40-60 minutes. SCP movement was determined by marking the location of a cell nucleus and using tools of the Metamorph program to record the position of the cell nucleus at the recorded time intervals. Average translocations of 10-25 individual cells in pixels/min were recorded, transformed to um/hr, and analyzed using Excel. For translocation of SCPs marked with Cell Tracker dye, the fluorescent SCPs in a microscope field were identified by capturing an image with fluorescence illumination to identify the SCP, and then fluorescent SCP migration along axons was recorded by phase contrast time-lapse microscopy at one image/min for 45-90 min. After identifying the fluorescent SCPs, a random sample of SCP was selected and their positions along axons were measured at five min intervals. The mean velocity of migration was calculated as µm/hr migrated.

### Block ephrin-A binding

In one approach to blocking ephrin-A2 signaling, EphA3-Fc was added to media of DRG cultures at 20 ug/ml, based on other papers and on our preliminary data (25). Control studies involved addition of similar concentrations of Fc or the non-function blocking anti-L1 monoclonal antibody 8D9, which controlled for general effects of binding exogenous proteins to axons.

### EphA4-blocking peptide

In another approach ephrin-A/EphA4 interactions was blocked with the KYL peptide that binds EphA4 and blocks activation by ephrin-A2. The KYL and control peptides were generously provided by Dr. EB Pasquale of the Sanford Burnham Institute, San Diego, according the sequences in Murai et al (25). Our preliminary results showed reduced rates of SCP migration along DRG axons in the presence of 25 ug/mlKYL peptide.

## Cell Transfection

Dr. Catherine Krull (University of Michigan) provided ephrin-A2-GFP plasmids and control GFP plasmids for transfection of isolated SCPs. Electroporation procedures, provided by Dr. Krull, were used to transfect SCPs.

## ACKNOWLEDGEMENTS

These experiments and analyses were conducted in the Letourneau lab between md-2005 and mid-2007. They have never been published or reported in any abstract or written form. The funds to conduct this research were provided by <u>National Institutes of</u> <u>Health Grant HD19950</u> and by grants from the Minnesota Medical Foundation.

Images of E13 DRGs and other DRG images were kindly provided by Dr. Andrea Ketchsek, Assistant Professor at Arcadia University, Glenside PA 19038.

## VIDEO LEGENDS

1. Video 1 shows SCPs migrating from an explant of an E7 peripheral nerve segment. The substratum is laminin-coated. The microscope objective used is 40X. Images were taken every 30 seconds.
2. Video 2 shows an individual E7 SCP migrating on laminin. The broad leading lamella spreads forward, and the tail is retracted. The microscope objective is 63X, and images were taken every 30 seconds.
3. Video 3 shows the same SCP as Video 2. The video was started after adding 20 ug/ml blebbistatin. Note that the front of the cell broadens widely, and the tail of the cell is no longer retracted. 63X
4. 4 shows N-cadherin beads picked up and transported back from the front of an SCP. Contact with beads does not cause retraction of the leading lamella. 63x.
5. 5 shows an SCP that picks up ephrin-A2-coated microbeads. The areas of the SCP leading margin that touch an ephrin-A2 microbead contract and shorten. 63X.
6. 6 shows an SCP that touch ephrin-A2 microbeads at its posterior tail. The tail rapidly contracts into the cell body after the bead adheres. 40X.
7. 7. SCPs migrating from an E7 DRG explanted on laminin. SCPs can be seen extending forward along axons and then contracting their posterior to move ahead. 20X.
8. 8. SCPs migrating from an E7 DRG explanted on laminin overnight in the presence of 20 ug/ml EphA4-Fc. Fewer SCPs have migrated from the explant compared to medium with a control Fc, and although SCPs show active extension of processes, actual translocation is less frequent than in control conditions. 20X.
9. 9. SCPs migrating from an E7 DRG explanted on laminin overnight in medium with high [Ca++] in the presence of 20 uM STI571, which inhibits abelson kinase signaling downstream of ephrin-A2. In medium with functional N-cadherin, SCPs show active motility but actual cell translocation is less without STI571. 40X.
10. 10. SCPs migrating from an E7 DRG explanted on laminin overnight in low [Ca++] medium in the presence of 20 uM STI571, which inhibits abelson kinase signaling downstream of ephrin-A2. In medium without functional N-cadherin, SCPs show active motility and active cell translocation. 40X.
11. 11. SCPs migrating from an E7 DRG explanted on an L1 substratum in high [Ca++] medium. SCPs migrate along the axons and do not adhere to L1 substratum. 20X.
12. 12. SCPs migrating from an E7 DRG explanted on an L1 substratum in high [Ca++] medium. SCPs migrate along the axons and do not adhere to L1 substratum. 40X.
13. 13. SCPs migrating from an E7 DRG explanted on an L1 substratum in low [Ca++] medium without functional N-cadherin. SCPs are motile but do not migrate well along the axons and do not adhere to L1 substratum. 40X.
14. 14. SCPs migrating from an E7 DRG explanted on an L1 substratum in high [Ca++] medium in the presence of Rho kinase inhibitor H1152. SCPs show motile activity, but they do not tranlocate along the axons. 40X.
15. 15. SCPs migrating from an E7 DRG explanted on an L1 substratum in high [Ca++] medium in the presence of EphA4-Fc, which blocks ephrin-A2 activation of Eph4. SCPs show motile activity but they do not migrate along the axons. 40X.
16. 16. A fluorescent E7 SCP migrates along axons of an E7 DRG. The fluorescent SCP starts at upper center of image and migrates toward the top of the video frame. 40X.
17. 17. A field of fluorescently marked E7 SCPs migrating along axons of an E7 DRG on an L1 substratum in medium with functional N-cadherin. The fluorescently marked SCPs can be identified from images in Figure 30, which were taken at the beginning and the end of the videorecording. 20X.
18. 18. A field of fluorescently marked E7 SCPs migrating along axons of an E13 DRG on a laminin substratum in medium with functional N-cadherin. The fluorescently marked SCPs can be identified from images in Figure 31, which were taken at the beginning and the end of the videorecording. Translocation of the E7 SCPs along 13 axons is slow. 40X.

